# Rapid thalamocortical network switching mediated by cortical synchronization underlies propofol-induced EEG signatures: a biophysical model

**DOI:** 10.1101/2022.02.17.480766

**Authors:** Austin E. Soplata, Elie Adam, Emery N. Brown, Patrick L. Purdon, Michelle M. McCarthy, Nancy Kopell

## Abstract

Propofol-mediated unconsciousness elicits strong alpha/low-beta and slow oscillations in the electroencephalogram (EEG) of patients. As anesthetic dose increases, the EEG signal changes in ways that give clues to the level of unconsciousness; the network mechanisms of these changes are only partially understood. Here, we construct a biophysical thalamocortical network involving brainstem influences that reproduces transitions in dynamics seen in the EEG involving the evolution of the power and frequency of alpha/low beta and slow rhythm, as well as their interactions.

Our model suggests propofol engages thalamic spindle and cortical sleep mechanisms to elicit persistent alpha/low-beta and slow rhythms, respectively. The thalamocortical network fluctuates between two mutually exclusive states on the timescale of seconds. One state is characterized by continuous alpha/low-beta frequency spiking in thalamus (C-state), while in the other, thalamic alpha spiking is interrupted by periods of co-occurring thalamic and cortical silence (I-state). In the I-state, alpha co-localizes to the peak of the slow; in the C-state, there is a variable relationship between an alpha/beta rhythm and the slow oscillation. The C-state predominates near loss of consciousness; with increasing dose, the proportion of time spent in the I-state increases, recapitulating EEG phenomenology. Cortical synchrony drives the switch to the I-state by changing the nature of the thalamocortical feedback. Brainstem influence on the strength of thalamocortical feedback mediates the amount of cortical synchrony. Our model implicates loss of low-beta, cortical synchrony, and coordinated thalamocortical silent periods as contributing to the unconscious state.

**New & Noteworthy:** GABAergic anesthetics induce alpha/low-beta and slow oscillations in the EEG, which interact in dose-dependent ways. We construct a thalamocortical model to investigate how these interdependent oscillations change with propofol dose. We find two dynamic states of thalamocortical coordination, which change on the timescale of seconds and dose-dependently mirror known changes in EEG. Thalamocortical feedback determines the oscillatory coupling and power seen in each state, and this is primarily driven by cortical synchrony and brainstem neuromodulation.

## Introduction

Unconsciousness mediated by GABAergic anesthetics, such as propofol and sevoflurane, is characterized by a presence of alpha/low-beta (8-20 Hz) and slow oscillations (0.5-2.0 Hz) in the electroencephalogram (EEG) (Brown, Purdon, and Van Dort 2011; Lewis et al. 2012; Purdon et al. 2013; Mukamel et al. 2014; Scheinin et al. 2018; Malekmohammadi et al. 2019; Stephen et al. 2020). As effect site concentration (dose) is increased, the EEG shows less low-beta, lower frequency alpha, less alpha power, and increased slow wave power (Purdon et al. 2013; Mukamel et al. 2014; Stephen et al. 2020). The alpha (8-14 Hz) and slow oscillations are interdependent and the level of interdependence was found to indicate the level of unconsciousness (Purdon et al. 2013). (Purdon et al. 2013; Mukamel et al. 2014; Gaskell et al. 2017; Stephen et al. 2020) highlighted two states they called “peak-max” and “trough-max”. These are two states in which the slow and alpha rhythms are coupled by phase-amplitude coupling (PAC). In the peak-max, the amplitude of the alpha rhythm is maximal during the peak of the slow rhythm; in trough-max, the maximal alpha amplitude appears in the trough of the slow. The trough-max state appears close to the loss of consciousness (LOC), whereas the peak-max state occurs with deeper levels of propofol. However, during most of the time spent under propofol, the EEG does not reflect either of these states. This is shown in Fig. 1.

**Figure 1.**
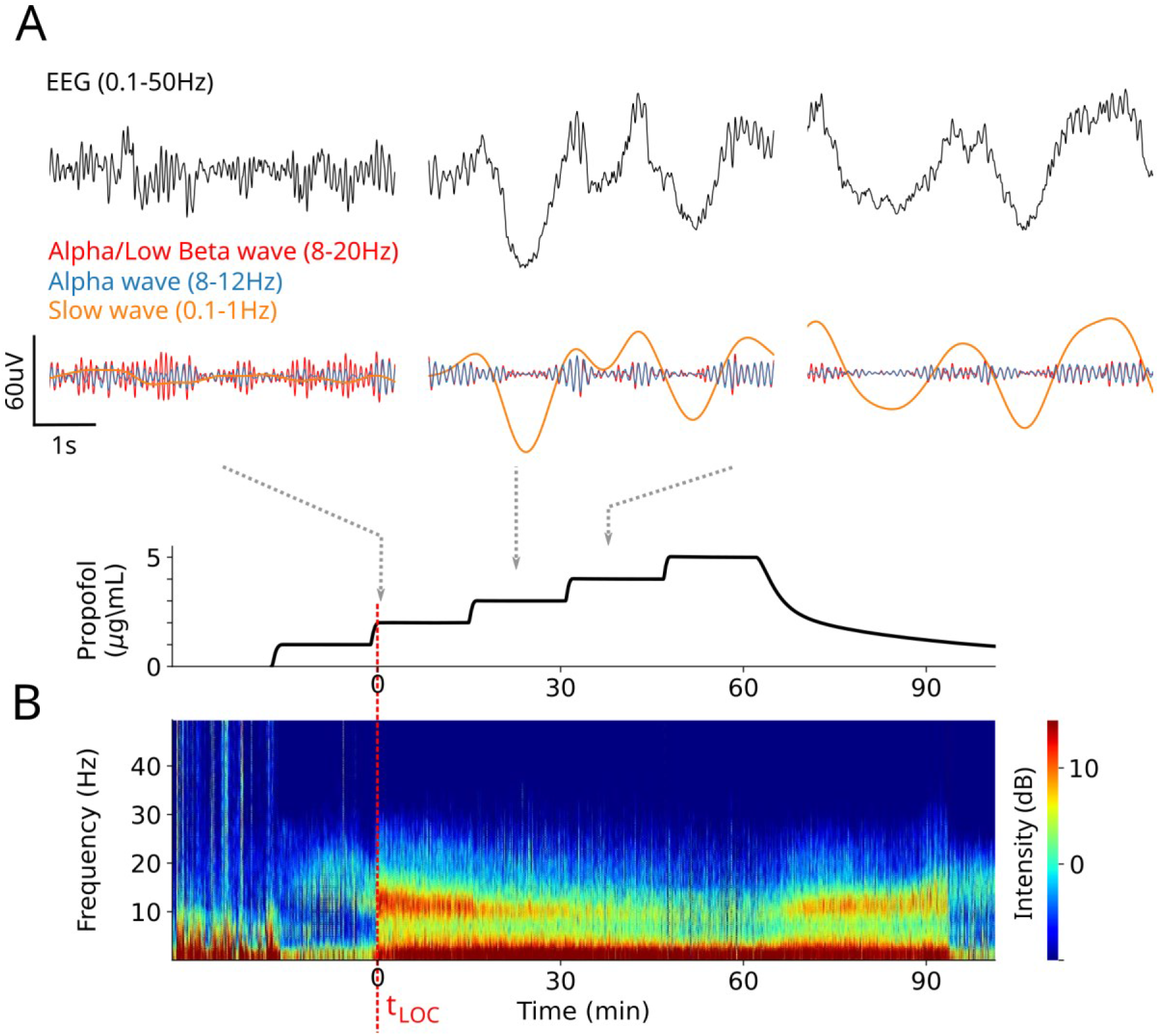
Typical spectral changes seen in the EEG as propofol dose is increased. Human experimental data from a single subject used in (Purdon et al. 2013; Mukamel et al. 2014) (see Methods). (A) Raw EEG traces (top) and EEG filtered at alpha/low-beta, alpha and slow (middle) at three different doses of propofol (bottom) starting near loss of consciousness (LOC) and up to but not including the state of “burst suppression” (Brown, Lydic, and Schiff 2010; Ching et al. 2012). Low-beta power appears predominately near LOC. (B) Spectrogram of the EEG during stepped increases in propofol dose in a healthy volunteer.

Anesthetics act on their molecular targets, and it is then through network mechanisms that they alter brain rhythms (Brown, Lydic, and Schiff 2010; Brown, Purdon, and Van Dort 2011). The network mechanisms underlying the EEG phenomena above remain unclear. Our previous work (Soplata et al. 2017) suggests alpha emerges from thalamus under propofol and can be modulated by an external source of slow oscillation such as that modeled in (Compte et al. 2003; Benita et al. 2012). Additionally, the alpha/slow peak-max increases as dose increases. However, our model was open loop, where cortex provides feedforward input to thalamus without feedback. It is unknown what dynamics are possible under a closed-loop interaction. Yet this interaction is essential during unconsciousness, for instance, to explain the anteriorization of alpha oscillations (Vijayan et al. 2013). In this work, we develop a biophysical model of closed-loop thalamocortical interaction that explains all the dose-dependent EEG phenomena above and shows the importance of thalamocortical feedback. Our Hodgkin-Huxley type model builds on (Soplata et al. 2017; Ching et al. 2010; Compte et al. 2003; Benita et al. 2012) to incorporate influences of cortex, thalamus and brainstem on EEG dynamics.

In this paper, we use 8-12 Hz as the range for alpha rather than the 8-14 Hz range used in (Purdon et al. 2013; Mukamel et al. 2014). We note that 8-12 Hz is the standard range for alpha in the cognitive literature (Van Diepen, Foxe, and Mazaheri 2019; Foster and Awh 2019). In the current paper, we aim to relate the propofol-induced rhythms to loss of cognitive function; and thus, we make a distinction between alpha (8-12 Hz) and low-beta (13-20 Hz).

We show that the dynamics on small space and time scales are highly complex: on each slow cycle, there is one of two network states, which can change after some indeterminate number of slow cycles. One of those states is called the “C-state” and is characterized by a continuous (“C”) alpha/low-beta oscillations in thalamus; the other state is called “I-state” in which there is thalamic spiking at an alpha frequency interrupted (“I”) by a time period within a slow oscillation cycle in which there is no thalamic activity. The I-state is highly related to what has been called peak-max, since the alpha activity is co-localized with the peak of the slow oscillation. The interaction of alpha, low-beta, and slow during the C-state is more variable than in the I-state, and the alpha portion includes what has previously been called “trough-max”. We show that the statistics of these states are dose-dependent, with higher doses of propofol corresponding to a larger percentage of the I-state. A significant finding of the work is that the statistics of the two states are strongly influenced by the synchrony of the cortical cells. Thus, the depth of anesthesia corresponds to the statistics of the I-states and C-states. Unconsciousness is associated with a prevalence of the I-state and thus a higher degree of cortical synchronization. Such increased synchronization has been reported experimentally (Gutiérrez et al. 2022). See the Discussion for more details.

Systemic administration of GABAergic anesthetics effects all structures in the brain, and the influence of these drugs on the brainstem can alter neuromodulatory systems (Moody et al. 2021). Our simulations show that when feedback from thalamus to cortex is potentiated because of these alterations, the synchronization of cortex is facilitated, and makes the switch to the I-state more probable. Endogenous noise in the system facilitates switching back to the C-state. Our work suggests that cortical synchronization due to potentiated thalamic feedback resulting from neuromodulatory alterations are key in understanding the mechanism of GABAergic anesthetics mediated unconsciousness. The loss of beta with increasing dose also has implications for loss of long-distance communication needed for consciousness.

## Results

### Model of propofol acts on cortical, thalamic and brainstem biophysics

We develop a model consisting of interacting thalamic and cortical circuits (see Methods). Our goal is to understand, and relate to loss of consciousness, the physiologic and network mechanisms related to EEG changes as propofol dose increases (Purdon et al. 2013; Mukamel et al. 2014; Flores et al. 2017; Stephen et al. 2020; Bastos et al. 2021) (Fig. 1). Among the spectral features that we investigate with our modeling are: (a) increased co-localization of the alpha with the peak of slow as dose is increased (Fig. 1A), (b) increased amplitude of the slow oscillation with increasing propofol dose (Fig. 1A), (c) increased low-beta power near loss of consciousness (LOC) and return of consciousness (ROC) (Fig. 1A,B), and (d) decreased mean frequency of alpha and decreased alpha power with higher doses (Fig. 1B). We show below that each of these features reflects properties of the model suggestive of mechanisms of loss of consciousness.

We model the cortical circuit using 100 pyramidal cells (PY) and 20 cortical interneurons (IN) from (Compte et al. 2003; Benita et al. 2012) (Fig. 2A). The pyramidal cells are modeled with two compartments: the soma (PYso) and the dendrite (PYdr). The thalamic circuit is modeled as in (Soplata et al. 2017), except with 20 thalamocortical cells (TC) and 20 thalamic reticular neurons (TRN). Our model currents are conductance-based with Hodgkin-Huxley dynamics. Details of the currents used in each neuron type, network connectivity, and all other aspects of the model can be found in the Methods and Appendix.

**Figure 2:**
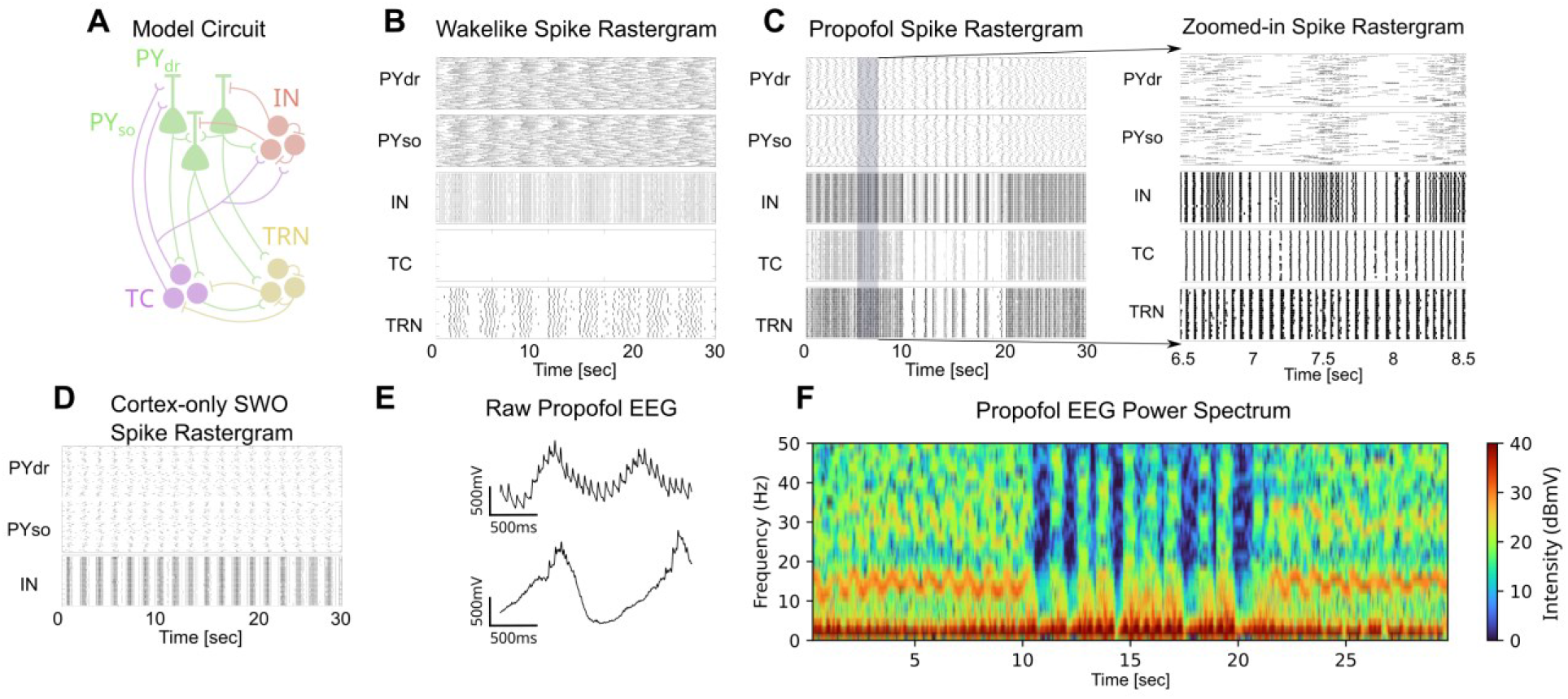
Propofol enables thalamocortical network slow and alpha oscillations. (A) Schematic of thalamocortical network model. TC: thalamocortical cells; TRN: thalamic reticular cells; PY: pyramidal cells (PYdr: dendrite, PYso: soma); IN: cortical interneurons. EEG is modeled as the sum of PYdr voltages. (B) Rastergram of each cell/compartment in wake-like mode with no sensory input, in which each black line represents a spike by each cell/compartment. (C) Rastergram of a simulation under propofol conditions. (Left) Simulation over 30 seconds. (Right) A zoom-in showing alpha/low-beta spiking in thalamus. (D) Rastergram of a cortex-only simulation showing slow wave generation in the absence of thalamus when the maximal conductance of the K(Na) current was 0.10 mS/cm^2^ representing the level seen during sleep. (E) Variations in model EEG. (Top) EEG taken from 23 to 25 seconds. Bottom EEG taken from 11 to 13 seconds. (F) Spectrogram of the model EEG from the simulation shown in (C).

We modeled the addition of propofol as five changes from wake-like conditions. The first is an increase in the maximal conductance and the inhibition time constant of the GABA receptors, which are known to be produced by propofol (McCarthy, Brown, and Kopell 2008; Ching et al. 2010). The second is a decrease in the maximal conductance of the TC cell H-current (Ying et al. 2006). Both were used in our previous work without feedback connections from thalamus to cortex (Soplata et al. 2017). The last three effects of propofol account for its action on brainstem circuits: specifically, propofol potentiates inhibitory GABAergic circuits in various arousal centers in brainstem (Brown, Purdon, and Van Dort 2011), including cholinergic centers. We focus on the enhanced inhibitory effects on brainstem cholinergic circuits (but see Discussion for other types of neuromodulation). We do not model the brainstem circuitry directly but rather model the effect of decreasing cortical cholinergic tone in the presence of propofol (Kikuchi et al. 1998; Nemoto et al. 2013; Meuret et al. 2000; Pal and Mashour 2021; Luo et al. 2020). This brainstem effect on decreased cortical cholinergic tone is modeled by strengthening (1) intracortical AMPAergic synaptic conductance, (2) TC→PY thalamocortical AMPAergic synaptic conductances, and (3) PY K(Na)-current maximal conductance (see Methods). As propofol dose is increased, we increase the strength of the intracortical and thalamocortical AMPA conductances. All other propofol changes are not further changed with dose. The EEG of the model is produced by the summation of the voltages in the dendrites of all cortical pyramidal cells (see Methods).

### Propofol generates slow waves in model cortex and alpha/low-beta waves in model thalamus

The “wake-like” condition without propofol produces a “depolarized relay state” (Destexhe et al. 1996) in thalamus as shown in Fig. 2B. In the depolarized relay state, the thalamus can transmit incoming signals; however, no such signals are included in the wake-like state and thus thalamus is silent (see (Ching et al. 2010; Krishnan et al. 2016) for similar models of wake-like thalamus). This simulation shows infra-slow oscillations in cortical cells (Fig. 2B), which are known to be present in the awake state (Krishnan, González, and Bazhenov 2018). The slow oscillation in our model originates from the cortex due to the K(Na) current (Compte et al. 2003) (see Methods). The firing rates of cells are consistent with known data (Dash, Autio, and Crandall 2022).

With propofol, the thalamocortical network produces prominent slow and alpha/low-beta oscillations that are visible in the raster plots (Fig. 1C). Here we show with a blow-up of Fig. 1C that the alpha/low-beta oscillation is visible in the thalamic TC cell spiking. The network dynamics can change in time in a way that will be examined in detail below. We find that the model cortex alone can generate slow oscillations and thus thalamic involvement is not needed (Fig. 2D). With propofol, an alpha oscillation arises in thalamus from the potentiation of GABA-A and the decrease in H-current, as shown in our previous work, which simulates thalamus in the absence of cortex. A description of the alpha generating mechanism is in (Soplata et al. 2017)in the section entitled “Propofol induces sustained alpha via changing the balance of excitation/inhibition.” This alpha uses the interaction between TC and TRN and engages thalamic spindling mechanisms. In our previous work, we did not include thalamocortical feedback; our current work shows that the thalamic alpha oscillations persist in the presence of thalamic feedback (Fig. 2C).

Our model EEG shows strong slow and alpha/low-beta oscillations (Fig. 2E). The alpha/low-beta oscillations are more evident (Fig. 2E, top) in the first and last 10 seconds of the simulation in Fig. 2C than during the middle 10 seconds (Fig. 2E, bottom). A spectrogram of the model EEG from the simulation in Fig. 2C, shows a continuous slow oscillation and a prominent higher frequency oscillation that switches between alpha/low-beta (8-20 Hz) and a lower average frequency alpha (∼ 8 Hz) (Fig. 2F).

### Propofol induces rapid switching between two distinct thalamocortical network states

With propofol, the network switches between two mutually exclusive network states on a rapid timescale (seconds) both on the single cell level and the population level (Fig. 2C, 3A-F), each of which can span over multiple cycles of the slow oscillation. The main difference between these states is the periodic cessation of spiking in the thalamus in one state but not the other; when this happens, the cortex and thalamus are simultaneously silent (Fig. 3A, D). The thalamic silence occurs because the thalamus enters a silent depolarized state, which stops the thalamus from spiking and thus thalamic input to cortex is lost. We call this network state the “I-state” (short for “interrupt”). In contrast, in the other network state, there are no thalamic silent periods because spiking from thalamus in continuous. Thus, we refer to this network state as the “C-state” (short for “continuous”).

**Figure 3:**
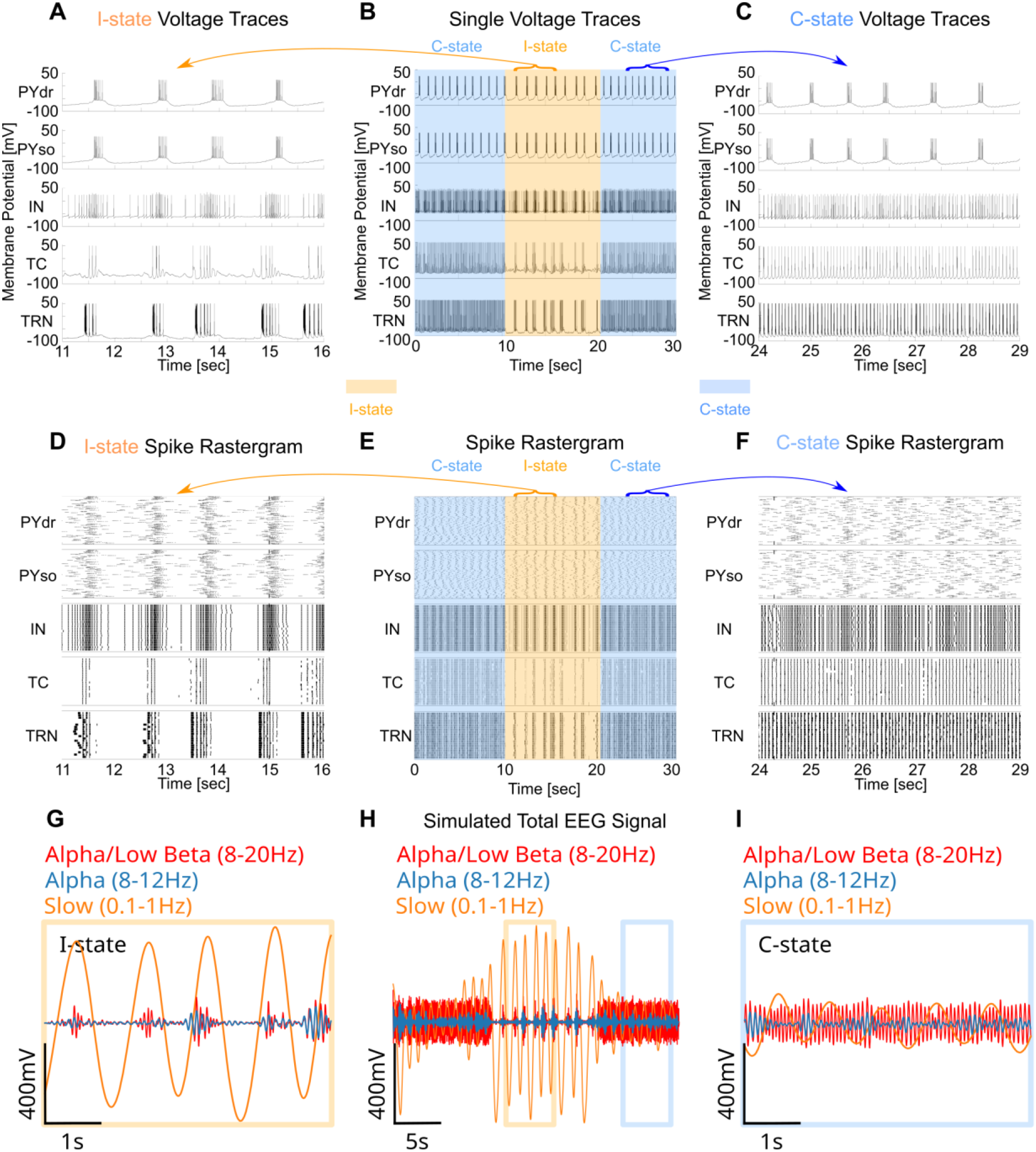
Different slow cycles display I-state or C-state during propofol. (A-C) Representative voltage traces of each cell/compartment (A) during an I-state, (B) across the entire propofol simulation, with I-state highlighted in orange and C-state highlighted in blue, and (C) during a C-state. (D-F) Rastergram of all spiking activity (D) during the I-state (E) across the entire simulation, with coupling regimes highlighted, and (F) during the C-state. (G-I) Model EEG filtered at alpha/low beta, alpha and slow (G) during the I-state, (H) across the entire simulation and (I) during the C-state.

Both states show slow oscillations and either alpha or alpha/low-beta oscillations in their spiking patterns (Fig. 3D-F) as well as in the model EEG (Fig. 3G-I) (See Methods for definition of model EEG). Although the alpha and low-beta rhythms are not a cortically generated rhythm, they appear in the model EEG due to the TC connections onto the pyramidal cell dendrites (Fig. 2A). During the I-state the thalamic fast frequency is predominately alpha (8-12 Hz) (Fig. 3D,G); during the C-state the fast thalamic frequencies range from 8-20 Hz and thus spans the alpha/low-beta range (Fig. 3F,I). These two network states correspond to different interactions between the slow and the alpha or alpha/low-beta in the EEG. In the I-state, there is always high amplitude alpha associated with the peak (or rising phase) of the slow oscillations (Fig. 3G); the trough of the slow oscillations corresponds to the simultaneous silent periods in thalamus and cortex (Fig. 3G). In contrast, the C-state has a variable relationship between alpha/low-beta and slow (Fig. 3I). The thalamic spiking persists continuously throughout the slow oscillation cycles with some variation in the relationship of the alpha/low-beta amplitude to the phase of the slow. In particular, the alpha and low-beta tend to couple at different phases of the slow oscillation during the C-state. Our simulations suggest that one possible coupling during the C-state is a transient coupling of alpha to the trough of the slow oscillation as seen on some slow cycles in Fig. 3I, as well as in experimental literature (Purdon et al. 2013; Mukamel et al. 2014). When this occurs, the low-beta couples to a different phase of the slow oscillation (Fig. 3I). Note that, in the C-state, the slow oscillation has lower amplitude than in the I-state (compare Fig. 3G and Fig. 3I).

In summary, the thalamocortical network rapidly switches between two states with propofol. The I-state is characterized by periods of coordinated thalamic and cortical silence as well as high cortical synchrony. In contrast, the C-state displays ongoing activity in thalamus and less cortical synchronization. The EEG during the I-state shows alpha oscillations but lacks low-beta oscillations, large amplitude slow oscillations, and co-localization of the alpha to the peak of the slow. In contrast, during the C-state the EEG has alpha and low-beta oscillations, smaller amplitude slow oscillations and variable alpha/slow coupling with alpha coupling to the trough being one possibility.

### Cortical synchrony determines thalamocortical state by modulating network feedback dynamics

Cortical synchrony is a critical determinant of thalamocortical state under propofol determining both the thalamic state (continuous spiking or interrupted spiking) as well as the spectral features of each state. The spiking in the cortex is noticeably more synchronized during the I-state than in C-state (Fig. 3D, F). The level of cortical synchrony affects the depolarization level of the thalamus and thus the thalamic spiking dynamics. Indeed, the less synchronized cortex in the C-state allows the thalamus to remain hyperpolarized and thus to continue engaging spindle dynamics (Soplata et al. 2017), while the additional synchronization in cortex during the I-state depolarizes the thalamus into its silent depolarized state. Thus, cortical synchrony is a critical determinant of thalamocortical state under propofol.

The C-state and I-state have different types of thalamocortical interactions under propofol. In the C-state, when the thalamus is spiking continuously (due to low cortical synchrony), the thalamus provides positive feedback to the cortex. In the I-state, the synchronous cortical input stops the thalamus by putting it its silent depolarized phase, whereas cortical silence promotes thalamic spiking (due to the hyperpolarization of the thalamus). Thus, the type of feedback in the I-state is homeostatic: cortical excitation leads to negative feedback from thalamus via silencing thalamus, whereas reduced cortical firing causes excitatory feedback from thalamus. The switch in feedback regimes (and thus network state) is abrupt because, as soon as the cortical synchronization level is sufficient to silence the thalamus, the network switches to homeostatic feedback.

The type of corticothalamic feedback dictates the spectral coupling observed during the different states. In the C-state, the alpha often occurs near the trough of the slow wave, while the low-beta occurs more towards the peak of the slow wave. This phenomenon stems from the influence of cortical spiking on the excitation level of the thalamus: when cortical spiking is low, as in the trough of the slow wave, the thalamus is more hyperpolarized and thus has a lower frequency of spiking (alpha spiking). In contrast, when the cortical spiking is higher, as in the peak of the slow wave, the thalamus is more depolarized and thus the thalamic spiking is at beta. The positive feedback from thalamus to cortex engages the activity-dependent K(Na)-current with the thalamus giving less excitation (alpha) to cortex when the cortical activity is low and more excitation (low-beta) when the cortical excitation is high. Note that this accounts for increasing frequency of the K(Na)-mediated slow oscillations between the wake-like state (∼ 0.2 Hz) and the C-state (1 Hz) (Fig. 2B, C). In contrast, when the network is in its homeostatic feedback regime (I-state), cortical synchrony leads to thalamic silence, which in turn leads to a profound loss of cortical spiking. This simultaneous loss of thalamic and cortical spiking is reflected in a large slow wave trough with no alpha coupling. The thalamus responds to the loss of cortical input by hyperpolarizing into its spindling regime and again spiking, now at predominately alpha, which depolarizes the cortex into its active phase. The diminished cortical K(Na), which decreased during the cortical inactive phase, additionally primes the cortex to spike more synchronously in response to the thalamic input. The more synchronous spiking during the cortical active phase is reflected in the EEG as a higher amplitude peak in the slow wave. The alpha spiking in the thalamus is coupled to the peak of the slow wave since this is the only phase at which the thalamus is active.

Summarizing the connection between cortical synchronization and thalamic state, we find that under propofol the thalamocortical network can abruptly switch between two dynamic networks states governed by the level of synchronization in the cortex. The spectral features of the EEG during these two states are the results of a switch in thalamic feedback dynamics: positive feedback during the C-state and homeostatic feedback during the I-state. In particular, we find that during the state of positive corticothalamic feedback (C-state), the amplitude and duration of the slow oscillation is controlled by the kinetics of the activity-dependent K(Na) current in the cortex, whereas during the state of homeostatic corticothalamic feedback (I-state), the amplitude and duration of the slow oscillation is influenced by the thalamic feedback. The larger amplitude slow waves during the I-state are a consequence of engaging the homeostatic thalamocortical feedback, while the low amplitude slow waves during the C-state result primarily from cortical K(Na) dynamics.

To verify the relationship between the cortical synchronization and the thalamocortical state, we tested whether we could change the thalamocortical state by introducing artificial cortical synchronization or de-synchronization. We applied 100 milliseconds of either a synchronizing or desynchronizing input to cortex during a period when the system was in a C-state. Artificial synchronization of cortex switched the thalamocortical network to an I-state, whereas with artificial the thalamocortical network remained in the C-state (Fig. 4A-E). These results support cortical synchronization as a driver of C-state to I-state transitions. When cortical synchronizing or de-synchronizing inputs were applied to the network when the system was in an I-state, we found that de-synchronizing inputs induced a transition to the C-state, whereas the network remained in the I-state in response to synchronizing inputs (Fig. 5). These results highlight cortical de-synchronization as a key determinant of I-state to C-state transitions, and thus a key determinant of thalamocortical state under propofol.

**Figure 4.**
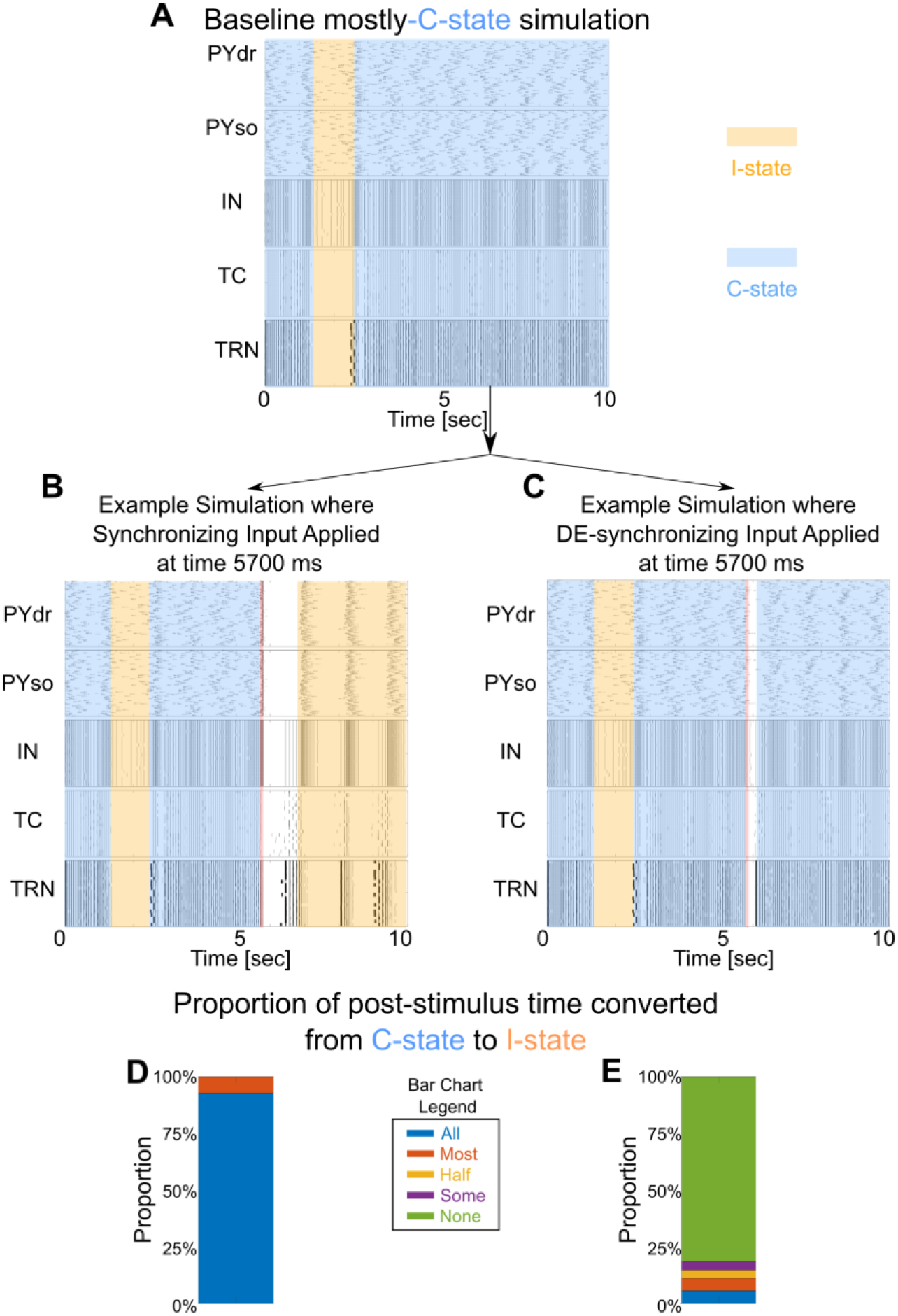
Cortical synchronization induces a C-state to I-state transition. (A) Rastergram of a “baseline” simulation in which most of the time is spent in the C-state. For every 100 millisecond interval expressing the C-state in the baseline simulation, two additional simulations were run: one simulation where a Synchronizing Input stimulus (see Methods) was applied to PYdrs during that specific 100 millisecond interval (example is shown in B), and another simulation where a DE-synchronizing Input stimulus (see Methods) was applied to PYdrs over that specific 100 millisecond interval (example is shown in C). (B) Rastergram showing an example simulation where Synchronizing Input was applied during the time 5700-5800 ms, causing the remainder of the simulation to exhibit I-state behavior. Red line marks the time of the input. (C) Rastergram showing an example of DE-synchronizing Input for the same time interval as B, but where the simulation almost completely remains in the C-state. (D) Bar graph showing, for all simulations receiving Synchronizing Inputs at different times, the proportion of post-stimulus simulation time that exhibited a C-to-I-state transition. Scoring was done by visual inspection, where each simulation was marked as “All” if >95% of simulation time after the Synchronizing Input exhibited I-state; analogously, each simulation was otherwise marked as “Most” = 95%-50%, “Half” = ∼50%, “Some” = 50%-10%, or “None” = <10%. The bar chart is the summation of all simulations receiving a Synchronizing Input and indicates that the artificial synchrony brought by the Synchronizing Input was very effective at converting C-states to I-states. (E) Similar to (D) but for baseline simulations receiving DE-synchronizing input, indicated that DE-synchronizing Input was not effective in eliciting C-state to I-state transitions.

**Figure 5.**
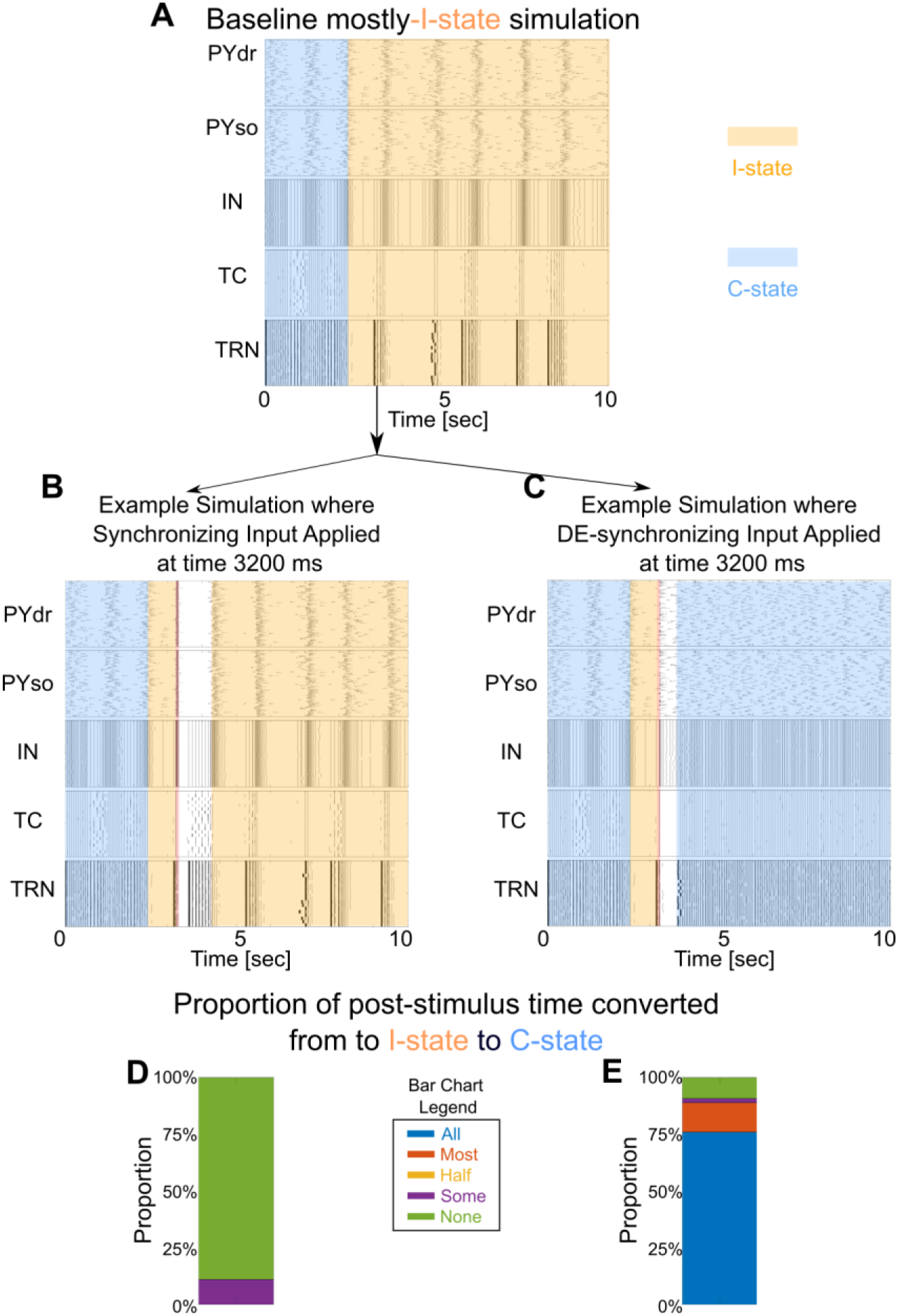
Cortical de-synchronization causes an I-state to C-state transition. This figure is analogous to Figure 4 but studying I-to-C transitions instead of C-to-I transitions. (A) Rastergram of a “baseline” simulation in which most of the time is spent in the I-state. For every 100 millisecond interval expressing the I-state in the baseline simulation, two additional simulations were run: one simulation where a Synchronizing Input stimulus (see Methods) was applied to PYdrs over that specific 100 millisecond interval (example is shown in B), and another simulation where a DE-synchronizing Input stimulus (see Methods) was applied to PYdrs over that specific 100 millisecond interval (example is shown in C). (B) Rastergram showing an example simulation where Synchronizing Input was applied during time 3200-3300 ms but did not lead to C-state behavior. Red line marks the time of the input. (C) Rastergram showing an example of DE-synchronizing Input for the same time interval as B, but where the remainder of the simulation exhibits C-state behavior. (D) Bar graph showing, for all simulations receiving Synchronizing Inputs at different times, the proportion of post-stimulus simulation time that exhibited an I-to-C state transition. Scoring was done as in Fig. 4. The bar chart is the summation of all simulations receiving a Synchronizing Input and indicates that the artificial synchrony brought by the Synchronizing Input was not effective at converting I-states to C-states. (E) Similar to (D) but for baseline simulations receiving DE-synchronizing input, indicated that DE-synchronizing Input was very effective in eliciting I-state to C-state transitions.

### Propofol potentiates TC feedback via brainstem neuromodulation, facilitating synchronization of cortex and a switch to the I-state as dose increases

In the previous section, we showed that cortical synchronization is a key driver of C-state to I-state transitions in corticothalamic circuits. Here we propose the physiological mechanism by which increased cortical synchronization occurs as propofol dose is increased.

We model increasing propofol dose by progressively decreasing ACh neuromodulation on corticothalamic circuits (see Methods): we increased maximal AMPA synaptic strength between cortical pyramidal cells 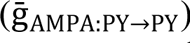 and from TC cells to cortical pyramidal cells 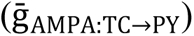, which strengthens feedback from thalamus to cortex. As propofol dose is increased in our model, the amount of time spent in the C-state progressively decreases and is replaced by more time spent in the I-state (Table 1). The progression in co-localization of alpha to the peak of the slow oscillation recapitulates the findings in subjects undergoing propofol-anesthesia induced unconsciousness.

**Table 1:** Proportion of time spent in the C-state when both TC→PY and PY→PY are set to the corresponding maximal gAMPA conductance. The units for gAMPA are mS/cm^2^. For each gAMPA value, measurements were taken across approximately 70 independent simulations that were each 30 seconds long.

| **gAMPA** | **Mean ± Std Dev** |
| --- | --- |
| **0.002** | 83.7% ± 9.81% |
| **0.003** | 83.26% ± 16.74% |
| **0.004** | 70.37% ± 25.87% |
| **0.005** | 44.39% ± 30.94% |
| **0.006** | 32.03% ± 34.22% |
| **0.007** | 15.61% ± 23.60% |
| **0.008** | 6.97% ± 15.61% |
| **0.009** | 4.15% ± 11.35% |
| **0.01** | 1.29% ± 3.26% |
| **0.012** | 0.65% ± 3.27% |

To determine whether the change to more I-states relies primary on TC→PY or PY→PY synapses, we looked at the incidence of I-states when changing one of these synapses at a time. We found that the dose-dependent increased time spent in the I-state is primarily due to increasing the TC→PY AMPA synapse (Table 2). The strengthened feedback leads to more synchronized cortical cells, which in turn causes the switch to the thalamocortical homeostatic feedback regime (see previous section) associated with I-state dynamics.

**Table 2:** Proportion of time spent in the C-state with differential values of TC→PY and PY→PY maximal gAMPA conductance. All values in form of “**mean** ± standard deviation”. For each set of gAMPA values, measurements were taken across approximately 40 independent simulations that were each 30 seconds long.

|  |  | TC→PY gAMPA (mS/cm<sup>2</sup>) |  |  |  |
| --- | --- | --- | --- | --- | --- |
|  |  | 0.004 | 0.006 | 0.008 | 0.01 |
| PY→PY gAMPA (mS/cm<sup>2</sup>) | 0.004 | 58.49% ± 25.07% | 32.69% ± 30.00% | 12.79% ± 14.31% | 4.82% ± 11.01% |
|  | 0.006 | 54.53% ± 32.70% | 21.33% ± 24.49% | 8.42% ± 10.33% | 3.15% ± 8.92% |
|  | 0.008 | 52.52% ± 25.46% | 27.92% ± 27.83% | 6.05% ± 11.04% | 4.67% ± 8.56% |
|  | 0.01 | 43.06% ± 25.29% | 17.71% ± 19.18% | 7.07% ± 13.46% | 2.36% ± 5.54% |

That the C-state to I-state switch requires thalamocortical feedback is supported by the finding that in the absence of TC feedback, the thalamocortical networks remain in the C-state (Fig. 6). It also shows that the cortical synchronization caused by the K(Na)-production of the slow wave does not produce sufficient cortical synchrony to induce the C-state to I-state transition. In contrast, we showed in the last section that the switch from the I-state to the C-state is facilitated by cortical desynchronization. This can occur physiologically under propofol due to cortical noise (Fig. 3).

**Figure 6.**
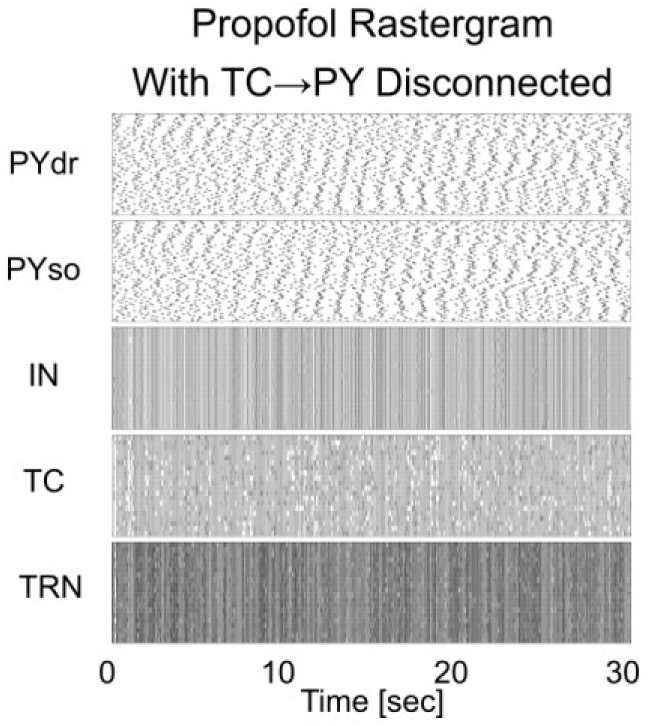
Lack of thalamocortical connections disables transitions from C-state to I-state. Rastergram of a “high-dose” 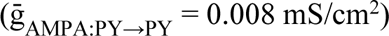 propofol simulation where TC→PY AMPA synapses have been set to zero. Simulations without thalamocortical feedback never exhibited an I-state.

The increasing time spent in the I-state with higher doses of propofol accounts for several dose-related EEG findings including: (1) less low-beta because this frequency is predominately in the C-state, (2) more lower frequency alpha because the alpha in the I-state has a lower average frequency than the alpha in the C-state (see Fig. 4B), (3) decreased alpha power because alpha is only present for a short period on the peak of the slow wave in the I-state, (4) loss of trough-max because trough-max is seen only in the C-state and only occasionally in that state, (5) increased peak-max due to thalamic alpha spiking occurring only during the peak of the slow wave in the

I-state, (6) increased slow wave amplitude/power due to the I-state having larger amplitude slow oscillations as a result of increased cortical synchronization and its coordinated cortico-thalamic silent states.

## Discussion

### Overview and Clinical Implications

The anesthetic propofol produces oscillatory signatures on the electroencephalogram (EEG): prominent alpha/low-beta oscillations (8-20 Hz), slow oscillations (0.5-2.0 Hz), and increased co-localization of the alpha to the peak of the slow with increasing effect site concentration (Mukamel et al. 2014). Here we use computational models to examine the role of brainstem, thalamus, and cortex in producing these oscillations and shaping the interactions between them. Specifically, our results help to understand the biophysical origin of the rhythms and the role of the rhythms in producing loss of consciousness.

One surprising finding in our simulations is that the thalamocortical network switches at the timescale of seconds between two dynamic states characterized by continuous alpha/low-beta in thalamus (C-state) or by transient interruption of the thalamic alpha (I-state). In the latter, the alpha is co-localized to the peak of the slow rhythm (“peak-max”), whereas in the former, there need not be consistent co-localization. What is called “trough-max” in the literature is here a special case of the C-state seen in the literature at sustained low doses of propofol that are used in experiments with volunteers; with clinical use, these stages are passed through quickly (Purdon et al. 2013; Mukamel et al. 2014; Stephen et al. 2020). In the I-state, cortical and thalamic silent periods are coordinated, which is the reason for the peak-max coupling. We further show that increases in propofol dose lead to statistically more prevalence of the I-state versus the C-state. Cortical synchrony is a key driver of the transitions between the I-state and the C-state: we show that corticothalamic state transitions are mediated by the level of cortical synchrony. We illustrate this by introducing artificial synchrony and desynchrony in cortical spiking during ongoing periods of the I-state or the C-state (Figs. 4 and 5). Cortical synchrony can switch the C-state to the I-state, and desynchrony can cause the I-state to switch to the C-state. One determinant of cortical synchrony in our network is thalamocortical feedback; this is strengthened by neuromodulatory changes related to propofol dose, biasing the expression of the I-state over the C-state as the dose of propofol increases.

Our model reproduces several clinical EEG observations as propofol dose increases: less low-beta power, lower frequency alpha, less alpha power, loss of trough-max coupling, increased peak-max coupling, and increased slow wave amplitude/power. The model provides physiological and network mechanisms for these effects, which give insights into how propofol works to produce loss of consciousness. The dynamic switch between two types of thalamocortical feedback, positive feedback in the C-state and homeostatic feedback in the I-state, determine the corticothalamic spiking patterns as well as the EEG spectral changes that correlate with each state. As dose is increased, brainstem effects on the thalamocortical feedback enable increased cortical synchrony, and thus, I-states become more prevalent. The increasing proportion of time spent in the I-state with larger doses of propofol in our model accounts for all the experimental spectral changes mentioned above associated with increasing propofol dose. Cortical synchronization has been found experimentally to relate to the level of unconsciousness (Gutiérrez et al. 2022), and thus these EEG measures may indicate the level of propofol-induced unconsciousness.

Our model helps explain why earlier work emphasized the trough-max state (Purdon et al. 2013; Mukamel et al. 2014): they were filtering in the alpha band (8-12 Hz), while our C-state shows non-slow oscillations between 8-20 Hz. As can be seen in Figs. 1 and 3I, the alpha band is sometimes found in the trough of the slow oscillation with beta oscillations at other phases. Lower doses of propofol associated with the time of loss of consciousness (LOC) are also characterized by a larger beta power than seen either at resting wakefulness or at higher doses of propofol (Purdon et al. 2013; Mukamel et al. 2014; Stephen et al. 2020; Krom et al. 2020; Malekmohammadi et al. 2019). Our modeling suggests this beta comes from thalamus in the presence of propofol. The model also suggests why the lower doses of propofol that commonly occur around LOC are associated with a light state of anesthesia (Mukamel et al. 2014): the beta oscillations communicated to the cortex are believed to be the basis of long-distance coordination in cortex (Siegel et al., 2012) and hence, signals from the thalamus can be transferred more widely throughout the cortex. By contrast, modeling has suggested that alpha disrupts such coordination (Weiner et al. 2023; Jones et al. 2000), which is consistent with alpha being the predominant rhythm with deeper levels of propofol anesthesia. Thus, a ratio of low-beta power to alpha power may be a way to monitor the depth of propofol-induced unconsciousness near the onset of LOC.

### Relation to prior modeling work

Our previous computational modeling (Soplata et al. 2017) suggested thalamus as the source of propofol-induced alpha (although not low-beta), and showed that dose-dependent coupling between alpha and slow occurred if slow oscillations were imposed on the thalamus from cortex via an “open-loop”. In this work, we simulate a “closed-loop” thalamocortical network of Hodgkin-Huxley cells with feedback from both thalamus to cortex and cortex to thalamus. Most of the results we find are a consequence of this feedback. Our cortical model is derived from models that can generate slow oscillations during sleep (Compte et al. 2003; Benita et al. 2012).

While prior modeling demonstrated thalamocortical circuits could produce propofol alpha oscillations (Ching et al. 2010; Vijayan et al. 2013), or propofol alpha and slow (Krishnan et al. 2016), our previous work (Soplata et al. 2017) was the first modeling investigation of which we are aware into the unique alpha-slow PAC dynamics of propofol. The essential results of the previous paper continue to hold in our current model: (1) The alpha is generated by propofol acting on thalamic circuits; (2) Our previous hypothesis that propofol-induced slow oscillations can come from the cortex is supported by the new modeling; (3) Cortical slow and alpha can interact in different ways depending on the hyperpolarization of the thalamus. However, details of the coupling between alpha and slow oscillation are altered from our previous work by the interaction between the thalamus and cortex, which was not explicitly modeled in that prior work.

Some mechanisms have been changed from our prior model and yield a better fit with experimental data: (1) in the prior thalamus-only model, the maximal thalamic spiking frequency under propofol was alpha. In this modeling work, the thalamus can spike as high as low-beta (∼ 20 Hz) (which has been seen in experiments (Ujma et al. 2022; Wang et al. 2022)). (2) in the previous model, thalamic alpha emerged only during the cortical silent phase of the slow during low-dose propofol and only during the cortical active phase of the slow during high-dose. In the current model, in which the hyperpolarization level of the thalamus is influenced by the cortex, we find that, during the C-state, the thalamus does not hyperpolarize enough to completely stop thalamic spiking during the cortical active phase; rather, the thalamus produces alpha/low-beta throughout all phases of the slow oscillation. As a result of this, alpha/low-beta is more prominent, and thus will show higher power, during low-dose time periods, which the C-state is more prevalent in (see Fig. 1B) (Purdon et al. 2013; Mukamel et al. 2014; Stephen et al. 2020). (3) The addition of thalamocortical feedback allows changes in cortical synchronization with increase in dose: continuous alpha/low-beta in thalamus results from less cortical synchronization, and hence the slow waves that appear in cortical EEG have a significantly lower amplitude and greater variability (Fig. 1B,C; Fig. 3 G,I) during the C-state than during the I-state (Mukamel et al. 2014). (4) Another difference from previous work is that, in the current model during the I-state, the thalamic silence during the cortical silent period is due to depolarization from corticothalamic excitation, rather than hyperpolarization as suggested by the thalamus-only model. This difference is significant in linking cortical synchronization to state changes: cortical synchronization depolarizes thalamus out of its alpha spiking into a silent state, changing the dynamics to the I-state. (5) In this paper, we explicitly model the cortex rather than using it just as an input to the thalamus. Thus, we were able to model the EEG by using the sum of pyramidal cell dendritic voltages (Kirschstein and Köhling 2009).

Other models of alpha oscillations exist in the literature with cortical as well as thalamic sources. Most of the modeling literature about cortical alpha concerns the awake state (Silva, Amitai, and Connors 1991; Jones et al. 2000; Bollimunta et al. 2011; Sigala et al. 2014; Lozano-Soldevilla, ter Huurne, and Oostenveld 2016; Köster and Gruber 2022) (but also see (Vijayan et al. 2013)). In the context of propofol anesthesia however, the prominent alpha observed in the EEG is likely to come from the thalamus: alpha oscillations under propofol are globally coherent, whereas slow oscillations are not (Lewis et al. 2012; Purdon et al. 2013). This suggests cortical alpha is coming from a non-cortical source such as the thalamus. Our earlier modeling suggests the thalamus can produce an ongoing alpha in the presence of propofol via potentiated GABA-A hijacking thalamic spindle mechanisms (Soplata et al. 2017). This is a prediction of both our previous and current model that remains to be tested experimentally. A recent experimental and computational study looking at oscillations with another GABAergic anesthetic, isoflurane, suggests this anesthetic can also generate thalamic alpha oscillations (Jiang et al. 2022).

### Propofol, slow oscillations, and brainstem-mediated neuromodulation

A notable interpretation of this model is that the anesthetic effects of propofol depend on neuromodulatory effects as much as on GABA-A inhibition and TC cell H-currents. ACh release in rodent frontal cortex can be decreased by 70-85% in the presence of propofol (Kikuchi et al. 1998; Nemoto et al. 2013). In the model, lowered ACh affects several cortical currents: (1) lowered ACh increases the K(Na) current (Schwindt, Spain, and Crill 1989; McCormick 1992) enabling slow oscillations; (2) lowered cortical ACh increases intracortical AMPAergic synaptic conductances (Compte et al. 2003; Benita et al. 2012; Krishnan et al. 2016); and (3) cortical muscarinic receptors respond to lowered ACh by increasing thalamocortical AMPA conductance (Kruglikov and Rudy 2008; Favero, Varghese, and Castro-Alamancos 2012). The latter is critical in our model for producing the greater synchrony of the cortex needed to switch to the I-state more often as dose is increased.

Noradrenergic tone is also decreased in the presence of GABAergic anesthetics: propofol and sevoflurane potentiate inhibitory GABAergic synaptic activity from the pre-optic area of the hypothalamus onto noradrenergic neurons of the locus coeruleus, which diffusely project to cortex among other areas (Brown, Pavone, and Naranjo 2018). In the cortex, norepinephrine (NE), as well as ACh, abolish slow oscillatory activity mediated by cortical Up and Down states (Favero, Varghese, and Castro-Alamancos 2012). Thus, lowered cortical NE due to propofol may work collaboratively with ACh in the producing increased slow oscillations in cortex.

Unlike ACh, which non-specifically suppresses PY→PY and TC→PY excitatory (AMPA), NE selectively suppresses intracortical excitatory inputs (Favero, Varghese, and Castro-Alamancos 2012). Although we find the dominant effect on cortical synchronization with propofol is due to increasing the thalamocortical AMPA conductance, NE likely contribute to the overall cortical synchronization by increasing the intracortical AMPA conductance.

We note that propofol likely also utilizes known slow mechanisms via its effects on non-cholinergic brainstem neuromodulatory systems. Propofol affects not just the cholinergic sources in the basal forebrain, laterodorsal tegmental area, and pedunculopontine tegmental area, but also inhibits the tuberomammillary nucleus, locus coeruleus, dorsal raphe nucleus, ventral periaqueductal gray, and lateral hypothalamus (Brown, Lydic, and Schiff 2010; Brown, Purdon, and Van Dort 2011). These areas respectively provide histamine, norepinephrine, serotonin, dopamine, and orexin/hypocretin to the cortex (Brown, Lydic, and Schiff 2010; Brown, Purdon, and Van Dort 2011). Many of these neuromodulators affect various potassium currents that are critical in known models of slow oscillations, including the K(Na)-current, the NaP-current, and potassium leak currents (Schwindt, Spain, and Crill 1989; McCormick 1992). These neuromodulators can also affect both excitatory and inhibitory currents in the cortex, and can change the relative impact of thalamocortical synapses (McCormick 1992; Favero, Varghese, and Castro-Alamancos 2012; Kuo and Dringenberg 2008). Additionally, cortical neuromodulation by NE, as well as ACh, is known abolish SWO and is active during awake/relay states (Moody et al. 2021). There is still much we do not understand about how all these neuromodulators work in concert together (Krishnan et al. 2016; Moody et al. 2021).

It is known that the cholinesterase inhibitor physostigmine reverses propofol LOC (Meuret et al. 2000); assuming LOC depends on slow oscillations and synchronization of the cortex, our model suggests that the engagement of ACh-modulated currents by propofol may explain this experimental result: physostigmine acts to increase ACh levels, which would decrease 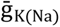 in our model and stop the K(Na)-dependent slow oscillation produced by propofol. ACh also acts to weaken thalamocortical connections (Favero, Varghese, and Castro-Alamancos 2012) and therefore release the cortex from over-synchronization; less synchronized states are associated with the awake state. Thus, loss of ACh may be a major contributor to propofol-induced loss of consciousness. The fact that our model requires neuromodulatory changes to produce propofol oscillations and their coupling suggests that the effects of propofol on the brainstem may be critical for its oscillatory phenomena, which is supported by active experimental research on propofol and other anesthetics (Moody et al. 2021; Minert, Yatziv, and Devor 2017; Minert, Baron, and Devor 2020; Muindi et al. 2016; Vlasov et al. 2021; Pal and Mashour 2021). Since the transition from C-state to I-state in our model is associated with lowering ACh (without changing GABAergic effects), we predict that a smaller dose of physostigmine may produce less I-state and more C-state and thus, may lead to more EEG low-beta, higher frequency alpha, increased alpha power, less peak-max, and lower slow wave amplitude.

### Propofol, slow oscillations, and sleep

Our work suggests that propofol utilizes not only thalamic spindling mechanisms (Soplata et al. 2017), but also natural sleep slow mechanisms and changes in neuromodulation to produce its oscillatory effects. The K(Na)-current is the primary mechanism of slow oscillation generation in the cortical sleep slow oscillation model we used (Sanchez-Vives, Nowak, and McCormick 2000; Schwindt, Spain, and Crill 1989; Compte et al. 2003). Most other slow oscillation models rely on a combination of changes to cortical excitatory/inhibitory plasticity and/or the persistent sodium current (NaP) (Bazhenov et al. 2002; Hill and Tononi, Giulio 2004; V. Crunelli et al. 2011; Sanchez-Vives and McCormick 2000; Timofeev et al. 2000; Krishnan et al. 2016). The NaP-current has been shown to be functionally coupled to the K(Na)-current (Hage and Salkoff 2012), and therefore the K(Na)-current may contribute to these mechanisms. Some models of slow UP state initiation, also called DOWN-to-UP transitions, rely on random cortical excitation (Timofeev et al. 2000), synaptic plasticity changes (Krishnan et al. 2016; Sanchez-Vives and McCormick 2000), or TC initiation of cortical slow UP states (V. Crunelli et al. 2011). In our simulations, the DOWN-to-UP transition is not reliant on the above mechanisms but rather initiates when the hyperpolarizing K(Na)-current in a PY cell has decayed sufficiently to allow significant PY spiking. In our I-state, however, the slow oscillation is primarily a manifestation of the change in thalamocortical feedback dynamics. The network feedback switches from a positive feedback dynamic in the C-state to a homeostatic feedback dynamic in the I-state. The homeostatic feedback produces wide swings in the amplitude of the slow oscillations, which differs from low amplitude K(Na)-mediated slow oscillation in the C-state. Future work designed to differentiate natural sleep slow oscillations versus general anesthetic slow mechanisms will enable finer-grained experiments into how the loss of consciousness occurs in these two distinct states.

### Propofol, memory consolidation, and aging

Our investigation of thalamocortical dynamics under propofol may have implications for memory and aging. During natural sleep, memory consolidation onto cortical axo-dendrite connections likely occurs during the nesting of hippocampal ripples during thalamic spindles, which themselves are nested inside thalamocortical sleep slow oscillations (Penagos, Varela, and Wilson 2017). Based on our current and previous work (Soplata et al. 2017), propofol alpha and slow oscillations hijack some of the same mechanisms used during sleep-related memory consolidation: propofol alpha derives from the thalamic spindling mechanism, and propofol slow oscillations engage the cortical K(Na) current also engaged by sleep. Proper memory consolidation requires correct encoding of worthwhile memories during sleep (Stickgold 2005), but if application of propofol abnormally activates some of the same oscillations in this process, this may lead to invalid memory consolidation or interfere with synaptic-dendrite networks involved in storing memory. A recent experiment showed promising results in using propofol to disrupt reconsolidation of traumatic memories (Galarza Vallejo et al. 2019), which could help treat Post-Traumatic Stress Disorder patients. Additionally, alpha power and, to a lesser extent, slow power under propofol may indicate a subject’s “brain age” (Purdon et al. 2015). Our modeling predicts that propofol alpha may predominately arise from the thalamus, and therefore a decrease in propofol alpha power across age could correlate with brain fitness via losses in the ability of the thalamus to burst at alpha or enter the spindling regime (Purdon et al. 2015), myelination retention of thalamocortical afferents (Peters 2002), or the strength of thalamocortical synapses onto cortical dendrites (Morrison and Baxter 2012).

### Caveats, Limitations, and Future Modeling

Our model is relatively small considering its significant complexity: there are a total of 160 neurons among multiple types in both cortex and thalamus. Larger models may sometimes display behavior that is not captured in smaller ones because of the degree of heterogeneity that is possible. Delineating such behavior is beyond the scope of this work. Also, not included in our model is complexity associated with the multiple layers of the cortex and the multiple kinds of nuclei in the thalamus. The use of Hodgkin-Huxley modeling in this study is justified when examining the effects of anesthetic drugs such as propofol, which work to change the kinetics of synaptic receptors and the conductances of intrinsic membrane currents that interact to sculpt the network behavior (Brown, Purdon, and Van Dort 2011). As such, the size and complexity of the model is driven by the questions asked. We found that simpler models, both in cellular components and neuron numbers, were not sufficient to replicate the spectrum of propofol-induced EEG changes explored in this paper and understand their mechanisms. As to further detail, our philosophy is to look for a minimal model, however complex, that will reproduce the interactions of the slow and alpha rhythms consistent with experimental results. Regarding the effects of brainstem modulation, we focused almost entirely on cholinergic modulation. Effects of other neuromodulators, briefly mentioned, are left for future work. Even the effects of ACh were not completely investigated; notably, we did not look at the effects of changing K(Na) strength with successively larger effect site concentrations of propofol.

Heterogeneity in TC→PY and PY→PY strength and connectivity across cortical regions and layers may contribute to diversity in cortical synchronization levels (Redinbaugh et al. 2020; Malekmohammadi et al. 2019) and therefore patient- or region-specific diversity in time spent in the I-state versus the C-state. Our simulations indicate that different effect site concentrations of propofol tend to express different proportions of these two network states on a small spatial scale. Our results also suggest that, under propofol, different local cortical networks may, on a fast timescale, switch between the I-state and the C-state even while a regional EEG signal predominantly shows a single type of dose-dependent PAC. By introducing region-specific heterogeneity to cortex (e.g., sensory and higher-order) and thalamus (e.g., core and matrix), future simulations may be able to investigate the significant spatiotemporal changes between low- and high-dose propofol. “Anteriorization” is a well-known phenomenon in which propofol administration initially leads to the loss of awake, occipital alpha and an increase in frontal alpha (Tinker, Sharbrough, and Michenfelder 1977; Cimenser et al. 2011; Vijayan et al. 2013). This frontal alpha is at its strongest and most persistent state during low-dose propofol, before spreading to become region-nonspecific during high-dose propofol (Cimenser et al. 2011; Purdon et al. 2013; Mukamel et al. 2014; Stephen et al. 2020) and decreasing in power with increasing dose (Gutiérrez et al. 2022). Slow power is also greater during high-dose than low-dose propofol (Purdon et al. 2013; Mukamel et al. 2014; Mhuircheartaigh et al. 2013; Lee et al. 2017) and may modulate higher frequencies more in frontal regions during high-dose (Stephen et al. 2020). We note that our current modeling captures many of these results for frontal cortex: (1) alpha is strongest and most persistent during low-dose propofol; (2), alpha decreases in power with increasing dose; (3) slow power is greater during high dose than low-dose propofol.

In the future, modeling multiple cortices will allow us to probe why holding propofol at a low-dose results in trough-max coupling that is most prevalent in frontal cortex (Mukamel et al. 2014), why there is stronger frontal slow modulation at higher doses (Stephen et al. 2020), and why there is increased thalamocortical alpha coherence in this region (Flores et al. 2017). Modeling multiple cortices will also allow us to explore coherence, phase (Malekmohammadi et al. 2019), and firing rate (Krom et al. 2020) discrepancies found between frontal and sensory regions under anesthesia. Understanding how region-specific heterogeneity affects cross-cortical communication and frontal cortex specifically may help to validate theories of loss of consciousness, including frontoparietal disconnection (Hudetz and Mashour 2016) and similar connectivity changes (Banks et al. 2020), brainstem changes to neuromodulation (Brown, Purdon, and Van Dort 2011), alpha blocking of processing (Palva and Palva 2007), and slow oscillation control of activity (Gemignani et al. 2015; Stephen et al. 2020).

There is strong recent evidence that thalamic simulation can reverse unconsciousness during low-dose propofol (Bastos et al. 2021). For our model to reproduce this, we would need to introduce additional mechanisms. One possible mechanism for this interesting phenomenon is that 30-seconds-long thalamic stimulation may activate group I metabotropic glutamatergic receptors (mGluRs) on the cortical cells receiving thalamocortical input. Thalamocortical activation of these receptors can produce depolarizing effects (Sherman 2014) and antagonism of cortical mGluRs leads to anesthesia-like effects (Suzuki and Larkum 2020). The very strong thalamocortical excitation elicited by thalamic stimulation may lead to activation of these mGluRs in the hyperpolarized cortical cells, depolarizing the cortical cells. This cortical depolarization could lead to a temporary reduction in cortical DOWN states (reducing SWO power) and an increase in higher power activity. Conversely, since mGluRs also exist at corticothalamic synapses (V. Crunelli et al. 2011; Sherman 2014), another contributor to the reversal of sedation could be that thalamic stimulation leads to thalamocortical excitation, which leads to corticothalamic stimulation that activates mGluRs, temporarily depolarizing the thalamus.

### Implications for unconsciousness

Higher doses of propofol corresponding to increased co-localization of alpha and the peak of slow have been associated with unarousable consciousness, whereas lower doses of propofol with less peak-max co-localization lead to a state of unconsciousness in which arousability is possible (Purdon et al. 2013; Mukamel et al. 2014; Scheinin et al. 2018). Our modeling suggests that the depth of anesthesia is associated with the degree of predominance of the I-state over the C-state. Since, in our model, cortical synchrony is a key driver of the transition from the C-state to the I-state and cortical desynchrony causes the opposite transition, our results suggest that increased cortical synchrony corresponds to deeper levels of unconsciousness. This has also been noted in the experimental paper (Bharioke et al. 2022) on unconsciousness and synchrony in layer 5 cortical cells. Such increased synchronization prevents the flexible and fast changes of coordination needed for normal cognitive processing (Cannon et al. 2014; Buzsáki 2006).

Our model suggests lower doses of propofol are associated with more time spent in the C-state. This state is associated with low-beta (as well as alpha) oscillations in the thalamus. As discussed above, beta oscillations have been documented to be highly involved in long-distance coordination in the brain (Siegel, Donner, and Engel 2012). Thus, the loss of this beta could contribute to the higher degree of unconsciousness associated with higher doses.

One unintuitive finding suggested by our model was that TC neurons may be depolarized into “relay mode” during I-state and could potentially relay sensory information during this window, even during deep anesthesia. In our simulations, strong corticothalamic excitation after synchronized active cortical states increased the membrane potential of TC cells during the I-state, as shown in Fig. 3A,D. This increase was enough to interrupt the intrinsic alpha bursts of the thalamus, but if this occurs at the same time as strong sensory input spikes, the TC cells may be depolarized enough to briefly relay sensory spiking information up to the cortex. Recently, even in humans under low-dose propofol anesthesia, auditory stimuli resulted in wake-like cortical neural activity in primary auditory cortex but not higher-order cortex (Krom et al. 2020). This suggests that some thalamic sensory relay may still occur under propofol anesthesia, even if changes to cross-cortical communication prevent its higher-order processing. Furthermore, in our simulations, the I-state may occur during individual slow cycles of both low- and high-dose propofol, indicating this brief sensory relay may occur at any point during propofol anesthesia (Malekmohammadi et al. 2019; Krom et al. 2020).

## Methods

### Model Design

Our Hodgkin-Huxley network, illustrated in Figure 2 A, consists of 100 cortical pyramidal dendritic compartments (PYdr), 100 corresponding cortical pyramidal somatic/axonal compartments (PYso), 20 cortical interneuron cells (INs), 20 thalamic thalamocortical cells (TCs), and 20 thalamic reticular neurons (TRNs). All equations and parameters used in the model are available in both the Appendix and the model code (Soplata 2023b; 2023a). The thalamic cells are identical to those used in (Soplata et al. 2017) and therefore derived from (Destexhe et al. 1996; Ching et al. 2010), except that we used a population size of 20 for each cell class rather than 50 due to memory/RAM limitations. The cortical compartments and cells are implemented according to their original description in (Compte et al. 2003), except that we include a simple leak-current in our PYdr compartments; we suspect that the original paper accidentally neglected to list this current. While there are many cortical slow models to choose from (Lytton, Destexhe, and Sejnowski 1996; Destexhe et al. 1996; Sanchez-Vives and McCormick 2000; Timofeev et al. 2000; Bazhenov et al. 2002; Destexhe and Sejnowski 2003; Hill and Tononi, Giulio 2004; Vincenzo Crunelli and Hughes 2010; V. Crunelli et al. 2011), we use this particular K(Na)-based sleep slow cortical model (Compte et al. 2003) due to its simplicity, experimental basis (Sanchez-Vives and McCormick 2000), and effective utilization in other slow models (Benita et al. 2012; Taxidis et al. 2013).

### Model Connectivity

All connections are illustrated in Figure 2 A and available in both the Appendix and the model mechanism code (Soplata 2023a). AMPA connections include from PYso to neighbor-only PYdr (PYso→PYdr also called PY→PY), from PYso to IN (PY→IN), from TC to TRN (TC→TRN), from TC to PYdr (TC→PY), from TC to IN (TC→IN), from PYso to TRN (PY→TRN), and from PYso to TC (PY→TC). Intracortical AMPA connections (PYso→PYdr and PYso→IN) included synaptic depression. NMDA connections include from PYso to PYdr and from PYso to IN and include synaptic depression. GABA-A connections include from IN to PYso (IN→PY), from IN to neighbor-only IN (IN→IN), from TRN to TC (TRN→TC), and from TRN to TRN (TRN→TRN). GABA-B connections are only from TRN to TC (TC→TRN). Finally, the only connections that are not chemical synapses are the simple compartmental connections between each PYdr compartment and its corresponding PYso compartment. Note that we use PY→PY to refer exclusively to AMPAergic PYso→PYdr connections.

For all synapses, each source cell is connected to its “nearest-neighbors” (2*radius+1) target cells, where the radius is 10 cells. This makes all connectivity ratios, or how many connections are projecting from each source cell, approximately 1:20. For our intrathalamic connectivity, since our population sizes are 20, this effectively makes all thalamic connections all-to-all connected, just like in (Ching et al. 2010; Soplata et al. 2017). For our intracortical connectivity ratio of 1:20, we based this primarily on the model from which we drew our SWO mechanism, (Compte et al. 2003) (1:20 with a standard deviation of 5), and subsequent work (Benita et al. 2012). Other models utilized either higher (Traub et al. 2005; Cruikshank et al. 2012) (ranging from 1:5 to 1:50, with most at 1:10 or 1:20) or lower (Krishnan et al. 2016) (1:10) connectivity ratios. Since the original model used a connectivity ratio that was comparable to similar models, and so as not to disturb network dependencies of the SWO mechanism, we elected to use their same intracortical connectivity ratio. Similarly, for the thalamocortical connectivity ratio of 1:20, we based our parameters on (Krishnan et al. 2016) (TC:PY ratio of 1:20). This gives us equal intracortical and thalamocortical connectivity ratios of both 1:20.

All synaptic conductances are normalized across the number of incoming connections of a given synapse type. For PY→PY connections, this is straightforward, since the size of the source and target cell populations are equal, leading to each PY cell receiving 20 of these connections. The maximal conductance of each of these incoming connections is computed to be 1/20th of the total maximal conductance for this synapse type. For connections where the source and population sizes are not equal, such as TC→PY connections, this is more complex. Each TC cell has 20 projections to the PY population, but since there are 20 TCs, this equals 400 total projections to go to 100 PY cells. Ultimately, this results in each PY cell receiving connections from 4 TC synapses, resulting in each synapse being normalized to 1/4th of the total maximum conductance. If the total maximum conductances of PY→PY and TC→PY are equal (see next paragraph), then each connection from a single source TC cell to a single target PY cell will have a larger maximal synaptic conductance than each single PY to PY connection, but there will be fewer TC to PY connections.

Except for the simulations done for Table 2, the total maximal conductances of our intracortical PY→PY and thalamocortical TC→PY AMPA synapses were kept equal to each other and changed simultaneously. We assumed that the ratio between these two maximal synaptic conductances could be held to be equal due to prior thalamocortical models using the same (Ching et al. 2010) or similar (Krishnan et al. 2016) (0.020:0.024) ratios. Additionally, by using equal values for these conductances, we could model the effects of propofol decreasing ACh on each of these synapse types in identical ways (see Table 1). We could then compare the effects on the system of treating these conductances heterogeneously (see Table 2) to understand their relative contribution to the case where they are set identically.

The total maximal conductances used for the PY→PY and TC→PY AMPA synapses ranged from 0.002 mS/cm^2^ to 0.012 mS/cm^2^. These values change in response to the concentration of ACh present, but the relationship between concentration and proportion of change in conductance is not clear, so we were required to explore a range of values; for the relationship of ACh to these conductances, see the subsection “Propofol effects” below. We initially based our intracortical and thalamocortical total maximal conductances on the default value used for PY→PY AMPA total maximal conductance in the original cortical SWO model paper (Compte et al. 2003): 5.4 nS, which is divided over a pyramidal dendritic surface area of 0.035 mm^2^ to give 0.0154 mS/cm^2^. Since this value is meant to correspond to an NREM sleep state which already exhibits low ACh concentration (Compte et al. 2003), we could use this value to simulate anesthetized states, but would need to decrease the value to simulate states of lower levels of anesthesia (see the aforementioned subsection below). We did simulate higher values such as 0.0154 mS/cm^2^ as in the original paper, but these were not significantly different than the I-state shown in the paper, and continued the trend shown in the Tables of a dominance of I-state.

### Model Inputs

For all simulations, in order to model background activity, excitatory Poisson spiketrains were input into all PY, TC, and TRN cells. These spiketrains had a firing rate of 40 Hz and were convolved with an exponential with a decay time of 2 milliseconds. The total maximal conductance of these virtual synapses were always held equal to that of the total maximal conductance of PY→PY and TC→PY AMPA synapses, except for the simulations in Table 2, where the Poisson inputs had conductances of 0.004 mS/cm^2^. See the mechanism code (Soplata 2023a) for implementation details.

For Figures 4 and 5, the “Synchronizing Input” was applied to all PYdr compartments for a duration of 100 milliseconds in each simulation and had a constant amplitude of 1.0 uA/cm^2^. For the “DE-synchronizing Input”, the input had a duration of 100 milliseconds, but each PYdr compartment received a different constant amplitude randomly pulled from a uniform distribution of -1.0 to +1.0 uA/cm^2^. In both cases, the amplitudes did not change for the duration of the stimulus and were 0 outside of the stimulus time.

### Propofol effects

Similarly to our previous work (Soplata et al. 2017), we model how increasing propofol affects the thalamus via changing three parameters: decreasing TC cell H-current maximal conductance 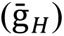, and potentiating all GABAA synapses via increasing maximal conductance 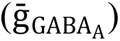 and GABAA decay time constants (τ_GABAA_). Propofol may decrease 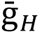 (Ying et al. 2006; Cacheaux et al. 2005), although the magnitude of this change is experimentally unknown (Chen 2005). To shift from a wake-like state to a propofol-anesthetized state, we decrease 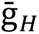 from 0.04 to 0.005 mS/cm^2^, which is in line with previous anesthetic and sleep research using this thalamic model (Destexhe et al. 1996; Vijayan et al. 2013; Ching et al. 2010; Soplata et al. 2017).

For our propofol simulations, we tripled 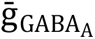 and τ_GABAA_ for all GABAA synapses (both cortical and thalamic), since doubling these GABAA parameters produced only I-state, but tripling lead to the presence of C-state. We originally based the magnitude of our propofol GABAA changes on prior modeling work (McCarthy, Brown, and Kopell 2008). In our previous paper (Soplata et al. 2017), we found that our thalamus-only network could produce persistent alpha oscillations if we doubled or tripled these GABAA parameters. In Figure 4 of (Soplata et al. 2017), we showed that thalamic persistent alpha occurred across a broader range of inputs when tripling the parameters compared to doubling. In the current paper, for all anesthetic simulation variations, doubling GABAA parameters produces little simulation time with persistent thalamic alpha oscillations. Instead, only by tripling GABAA parameters do the simulations produce C-state for a substantial or majority of simulation time. This may be due to effects on lower-dose behavior caused by additional cortical cell types that we did not include, namely from those of (McCarthy, Brown, and Kopell 2008).

Propofol decreases cortical acetylcholine (ACh) (see Introduction), and we model these cholinergic changes via increasing intracortical AMPAergic synaptic conductances 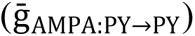 (Compte et al. 2003; Benita et al. 2012; Krishnan et al. 2016), TC→PY thalamocortical AMPAergic synaptic conductances 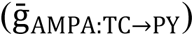 (Kruglikov and Rudy 2008; Favero, Varghese, and Castro-Alamancos 2012), and K(Na)-current maximal conductance 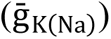 (Compte et al.

2003; Benita et al. 2012). ACh affects thalamocortical afferent synapses in different ways: decreased nicotinic ACh receptor activation weakens thalamocortical synapses, but decreased muscarinic ACh receptor activation strengthens them (Kruglikov and Rudy 2008; Favero, Varghese, and Castro-Alamancos 2012; Gil, Connors, and Amitai 1997; Hsieh, Cruikshank, and Metherate 2000; Oldford and Castro-Alamancos 2003; Eggermann and Feldmeyer 2009). Based on the rapid desensitization of nicotinic ACh receptors (Quick and Lester 2002), the slowly-changing, metabotropic nature of muscarinic receptors, and their similar shifts in natural sleep (McCormick 1992), we believe that muscarinic receptors could exert a stronger effect than nicotinic receptors on thalamocortical afferents, therefore increasing 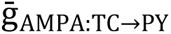 with increasing propofol dose.

The actual proportion of change that ACh can cause in the intracortical and thalamocortical synapses is unclear. Different computational models use a range of proportional increases to 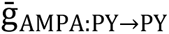 caused by ACh, including up to +75% (Vijayan and Kopell 2012) or +15% to

+100% (Krishnan et al. 2016). Data on how much ACh may increase 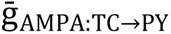 is much more scarce, but the increase may be as high as +300% (see Figure 8 F of (Favero, Varghese, and Castro-Alamancos 2012)). We therefore focused our synaptic changes on a wide range of decreases from 0.0154 mS/cm^2^, the derived value of the low-ACh sleep state used in the cortical SWO model of (Compte et al. 2003).

For the K(Na)-current maximal conductance 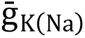, we only used two values: 0.10 mS/cm^2^ in our wake-like simulation (similar to the tonic, wake-like state of Figure 14 of the original paper (Compte et al. 2003), or 1.33 mS/cm^2^ for all anesthetic states. We specifically chose to not make small changes to 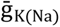 between anesthetic states because it has impacts on the frequency of the SWO produced, which the original paper investigates. Keeping the SWO frequency stable allowed us to more effectively investigate how PAC can arise, and how it can be changed by covariation of intracortical and thalamocortical synaptic strength.

### Specific simulation parameters

The changes to parameters for the wake-like simulation from those of the Appendix are as follows (all units in mS/cm^2^): 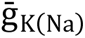, and all Poisson inputs set to 0.004, PY 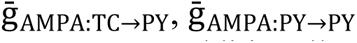. Additionally for the wake-like simulation, the following synapses were decreased by 60% compared to their values in the Appendix: 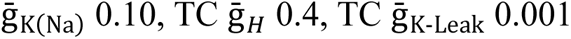. Our wake-like simulation was the only simulation where we included changes to the six synapse types in the previous sentence; this was done to better align with the wake-like simulations in Figures 14 and 15 of the original paper (Compte et al. 2003). The script used to run the wake-like simulation is available with the rest of the simulation scripts (Soplata 2023b).

For anesthetic states, all parameters were as given in the Appendix unless otherwise indicated, such as changes to 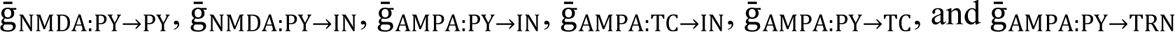 and 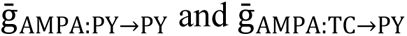 for Tables 1 and 2 and Fig. 6. All parameters are available in the mechanism (Soplata 2023a) and simulation (Soplata 2023b) code.

### EEG Model

We define our simulated EEG as the sum of dendritic voltages of PY cells as these are considered to be the main contributors (Kirschstein and Köhling 2009), bandpass filtered between 0.1-50 Hz. The results are unchanged if we considered PY somatic voltages.

### Simulations and Reproducibility

All of the simulation parameters (Soplata 2023b) and model mechanism code (Soplata 2023a) needed to reproduce the simulations shown in this work are available online on GitHub. All simulations were run using the “dev” branch of the MATLAB simulation toolbox DynaSim (Sherfey et al. 2018) located online. Individual simulations should be reproducible on a modern desktop computer with access to RAM of 32 gigabytes or higher.

Raw data for Tables 1 and 2 and the bar charts in Figures 4 and 5 were obtained by visual inspection from simulations which are reproducible from the simulation code (Soplata 2023b). The aggregate simulation data is available at Supplemental Data 1.

### Human Data for Figure 1

Human experimental data used in Figure 1 is from a single subject used in (Purdon et al. 2013; Mukamel et al. 2014). Analyses in Fig. 1A follows those in (Purdon et al. 2013; Mukamel et al. 2014). EEG traces were band-pass filtered to 0.1-1 Hz, 8-12 Hz and 8-20Hz using a Butterworth filter of order 2. EEG spectrogram in Figure 1B (not previously published) was computed using the multi-tapered method (Babadi and Brown 2014).

## Acknowledgements

Current affiliation for Austin Soplata is École Polytechnique Fédérale de Lausanne, Blue Brain Project, Campus Biotech, Chemin des Mines 9, 1202 Geneva, Switzerland. We thank the reviewers for their helpful comments on a previous draft of this paper. We would also like to thank Jason Sherfey, Erik A. Roberts, Emily Stephen, and Caroline Moore-Kochlacs for their suggestions during the investigation.

## Grants

All authors were supported by NIH Grant P01GM118269. Other sources include Guggenheim Fellowship in Applied Mathematics, NIH R01-GM104948, funds from MGH, and NSF DMS-1042134-5.

### Disclosures

Massachusetts General Hospital has licensed intellectual property for EEG monitoring developed by Drs. Brown and Purdon to Masimo Corporation. Drs. Purdon and Brown have a financial interest in PASCALL Systems, Inc., a company developing closed loop physiological control systems for anesthesiology. Dr. Purdon and Dr. Brown’s interests were reviewed and are managed by Massachusetts General Hospital and Mass General Brigham in accordance with their conflict-of-interest policies.

**Supplemental Data 1:**

**Simulation Data and Plots, available at** https://doi.org/10.6084/m9.figshare.22228858.v1

## Appendix: Equations for Computational Models

### 1 General Notes

Units of maximal conductances and currents for all equations below are in 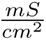 and 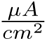, respectively, based on the original formulation of [Hodgkin and Huxley, 1952]. Capacitance *C_m_* for all cells was 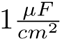. Thalamic circuitry is based on [Destexhe et al., 1993], [Destexhe et al., 1996], and [Ching et al., 2010] and is mostly identical to [Soplata et al., 2017]. Cortical circuitry is based on a from-scratch implementation of [Compte et al., 2003] and [Benita et al., 2012]. Thalamo-cortical and corticothalamic synaptic equations were derived from the same AMPAergic synapse equations as in [Soplata et al., 2017]. Occasionally, multiplication dots will be used (*·*) to help legibility. All simulations were run for 30 seconds of simulation time (unless otherwise indicated) and solved using Euler integration, with a time resolution (dt) of 0.01 ms.

### 2 (PYdr) Cortical Pyramidal Dendritic Cell Compartment Equations

#### 2.1 Voltage / Membrane Potential

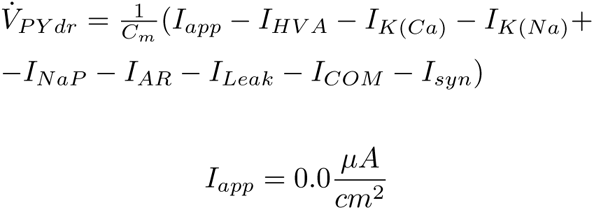

#### 2.2 (HVA) High-Voltage-Activated Ca Channel

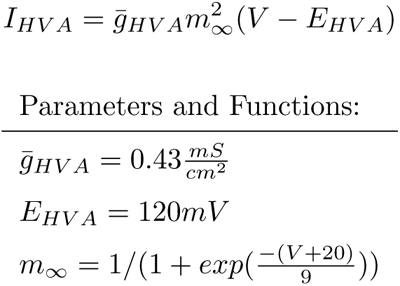

#### 2.3 (K(Ca)) Slow Calcium-activated K Channel

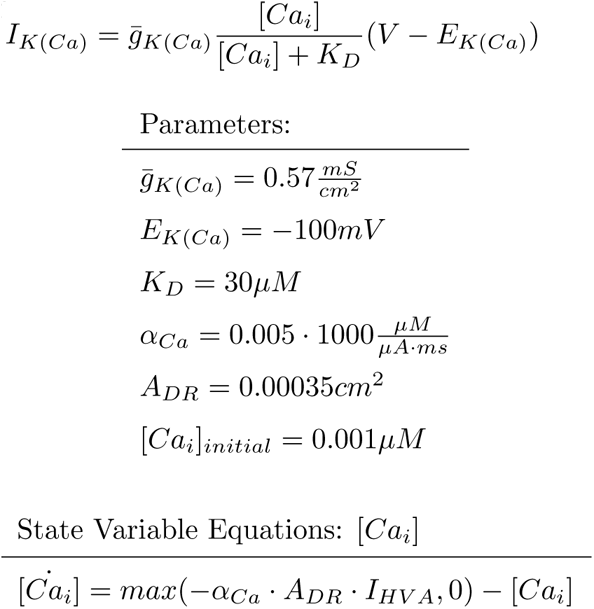

#### 2.4 (K(Na)) Na-activated K Channel

See the section “Non-synaptic Connection Equations”.

#### 2.5 (NaP) Persistent Na Channel

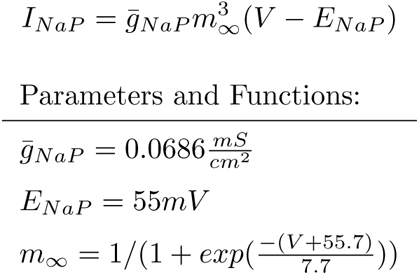

#### 2.6 (AR) Inwardly Rectifying K Channel

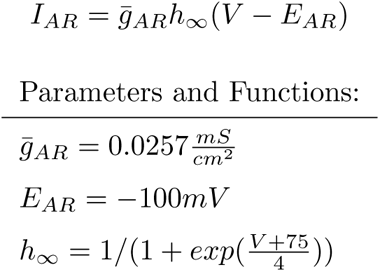

#### 2.7 (Leak) Leak Channel

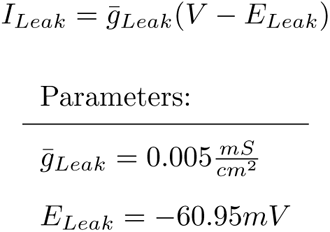

### 3 (PYso) Cortical Pyramidal Axosomatic Cell Compartment Equations

#### 3.1 Voltage / Membrane Potential

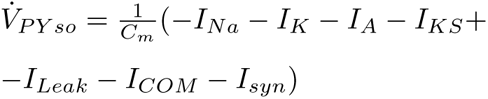

#### 3.2 (Na) Na Channel

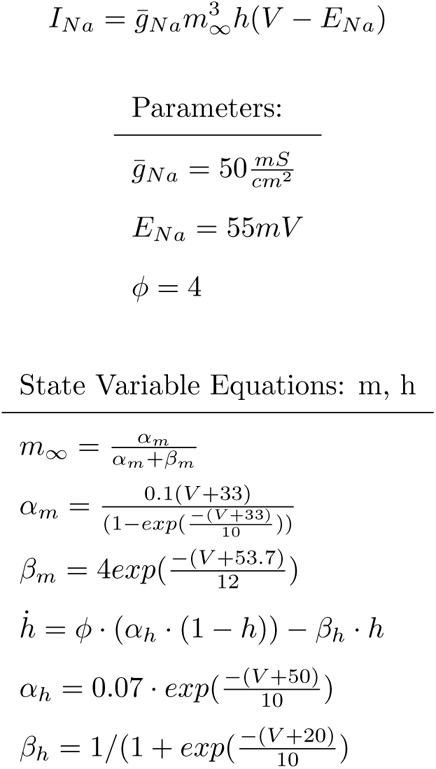

#### 3.3 (K) K Channel

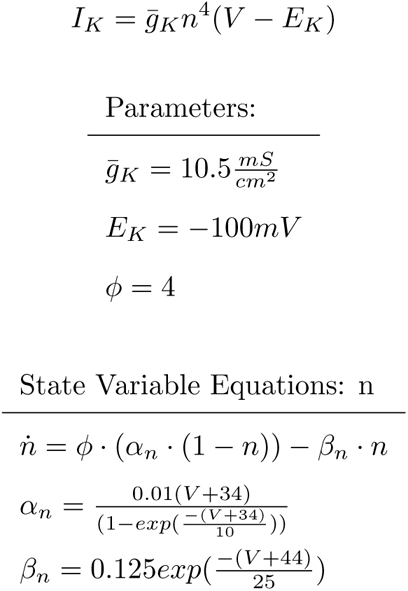

#### 3.4 (A) Fast A-type K Channel

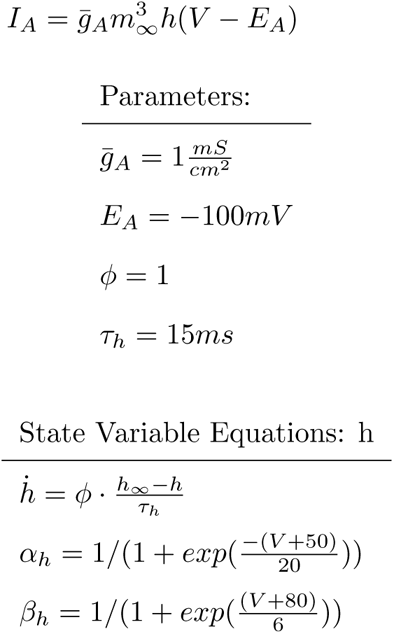

#### 3.5 (KS) Non-inactivating K Channel

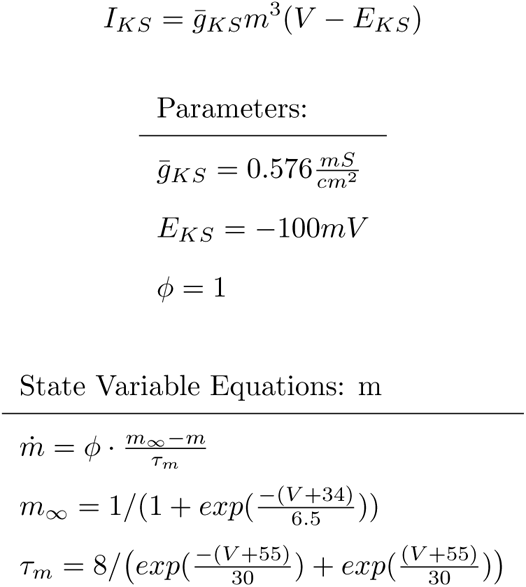

#### 3.6 (Leak) Leak Channel

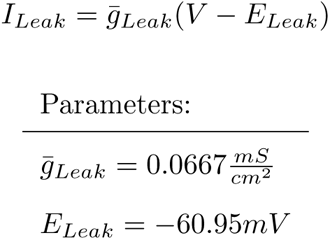

### 4 (IN) Cortical Interneuron Cell Equations

#### 4.1 Voltage / Membrane Potential

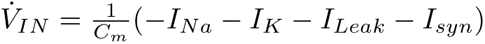

#### 4.2 (Na) Na Channel

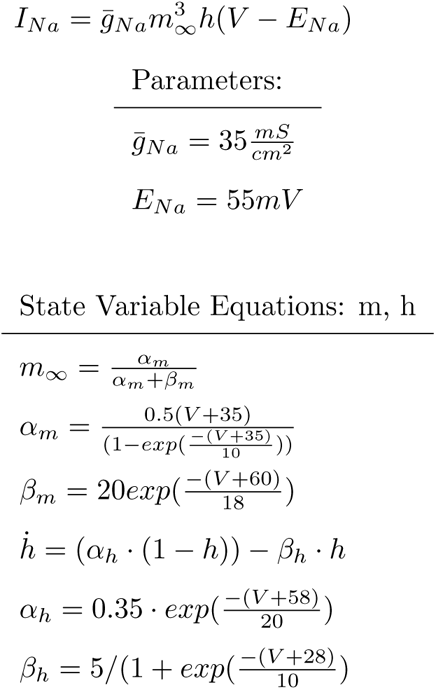

#### 4.3 (K) K Channel

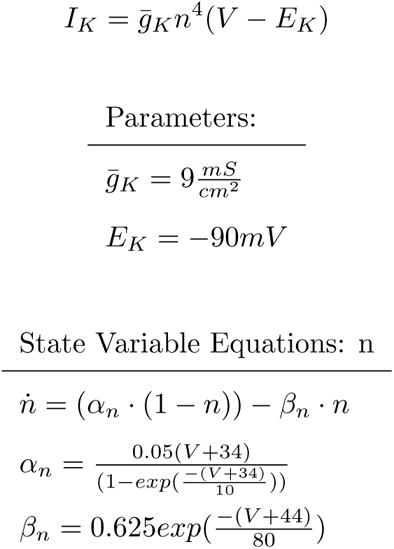

#### 4.4 (Leak) Leak Channel

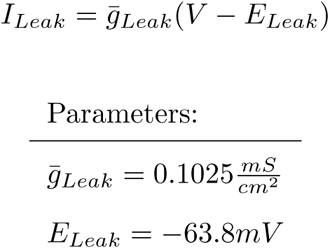

### 5 (TC) Thalamocortical Cell Equations

#### 5.1 Voltage / Membrane Potential

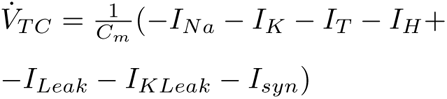

#### 5.2 (Na) Na Channel

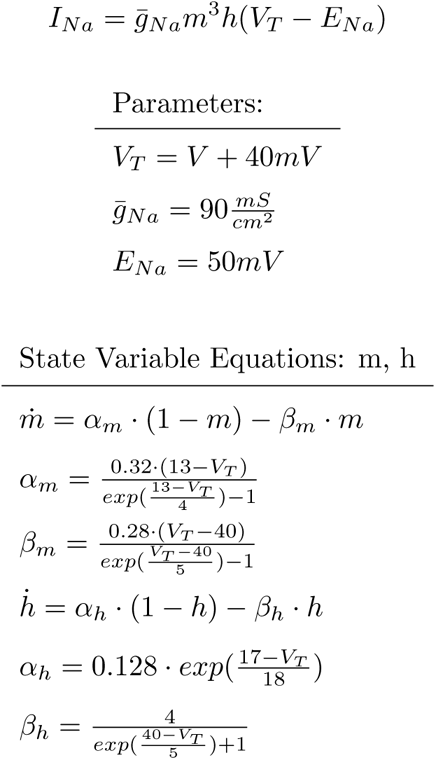

#### 5.3 (K) K Channel

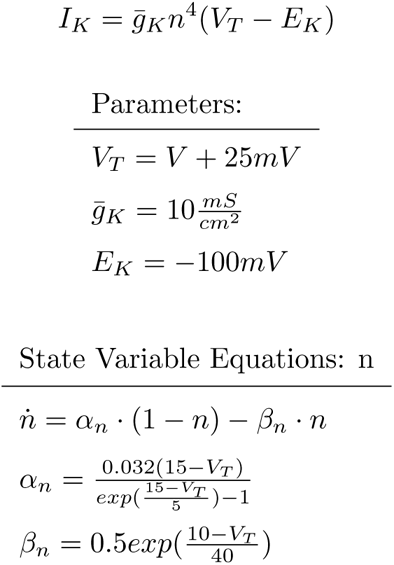

#### 5.4 (T) T-type Calcium Channel

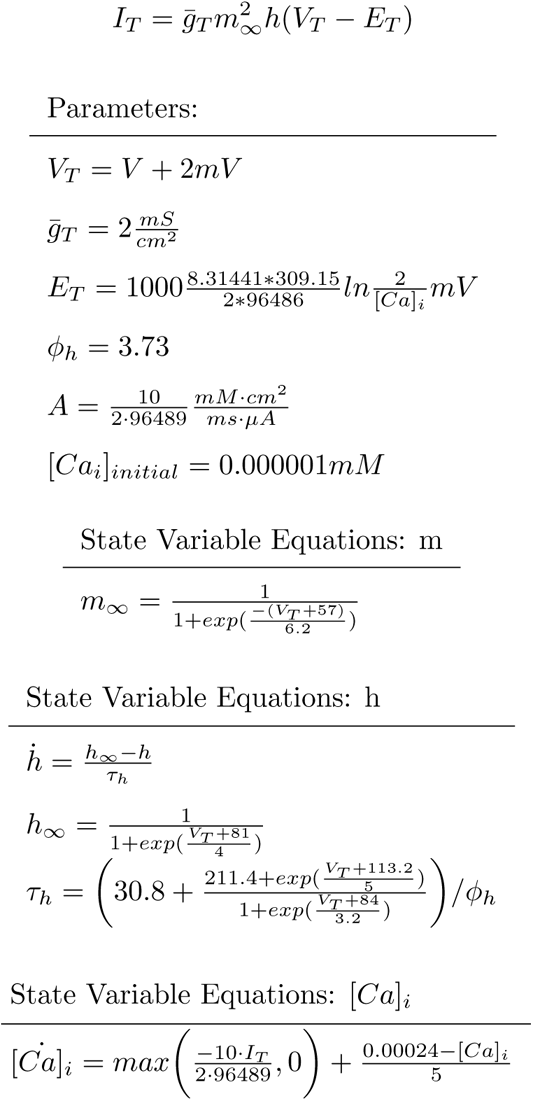

#### 5.5 (H) Hyperpolarization-activated Cation Channel

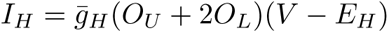

Note: This is the more complex [Destexhe et al., 1996] formulation of the H-current, not that of [Destexhe et al., 1993]. *O_U_*is the proportion of unlocked channels, *O_L_* is the proportion of locked-open channels, and *P*_1_ is the proportion of utilized substrate for channel locking.

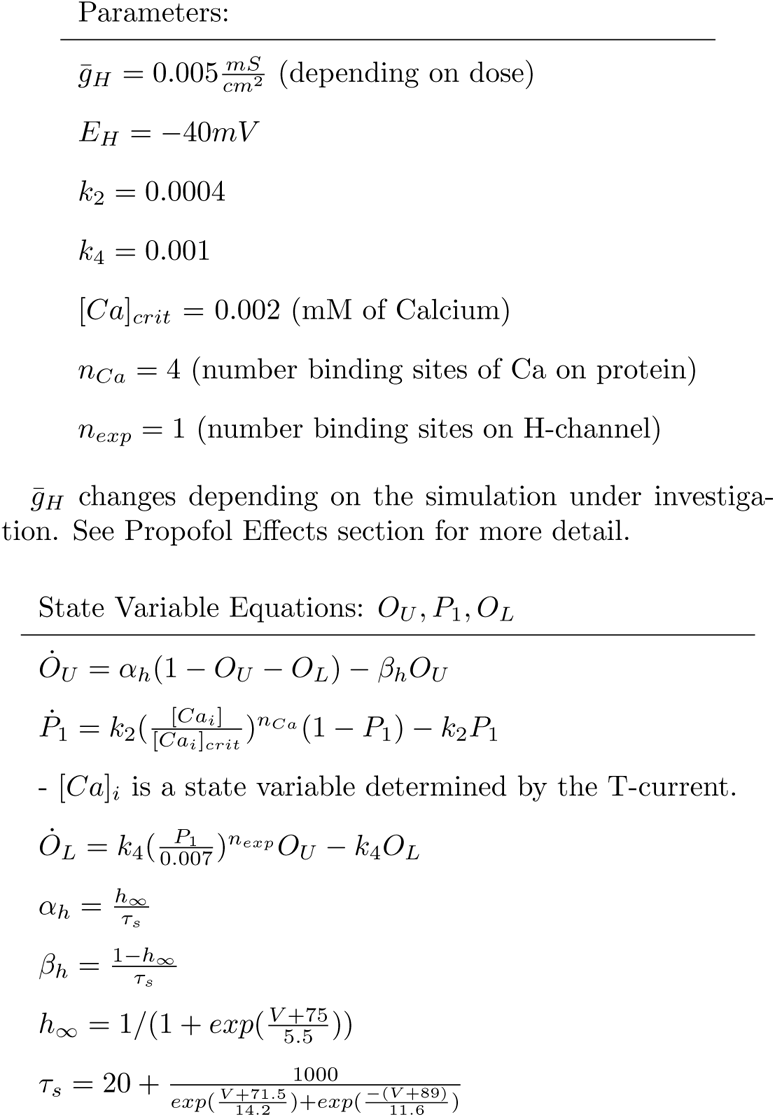

#### 5.6 (Leak) Leak Current

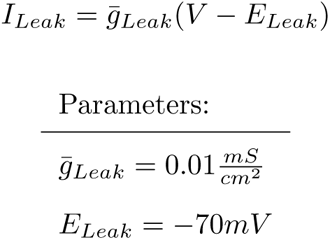

#### 5.7 (KLeak) K Leak Current

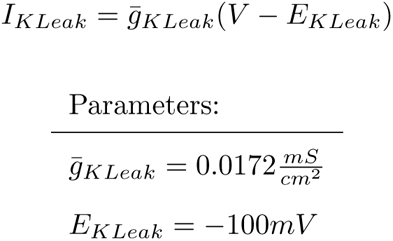

### 6 (TRN) Thalamic Reticular Nucleus Cell Equations

#### 6.1 Voltage / Membrane Potential

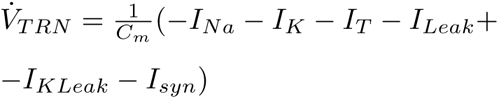

#### 6.2 (Na) Na Channel

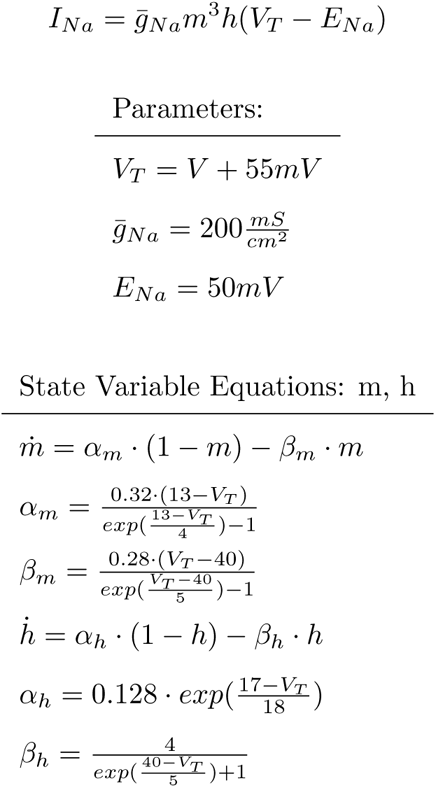

#### 6.3 (K) K Channel

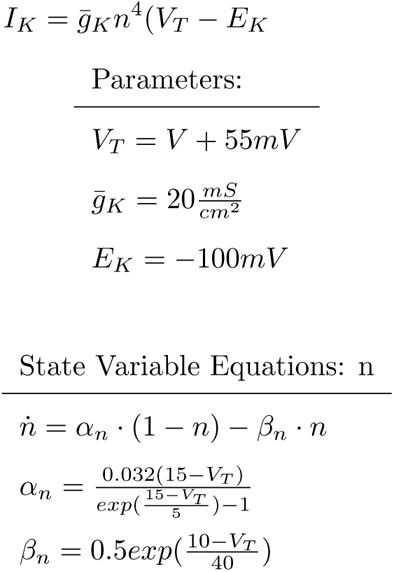

#### 6.4 (T) T-type Calcium Channel

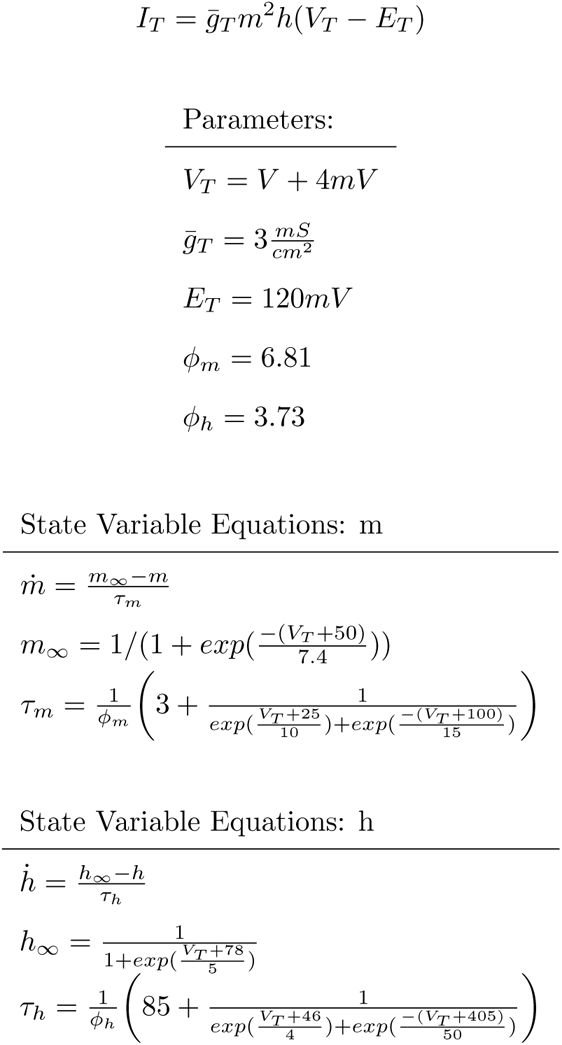

#### 6.5 (Leak) Leak Currents

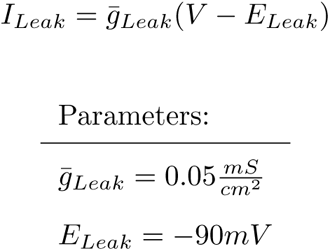

### 7 Non-synaptic Connection Equations

#### 7.1 Direct Compartmental Connections

Each PYdr compartment has a special connection to and from a single PYso compartment called *I_COM_*, meant to simulate voltage fluctuations between the axo-soma and dendrite of a single cell. Note that these conductances are not exact inverses of each other due to the difference in size between the two compartments. These two currents are calculated using the following:

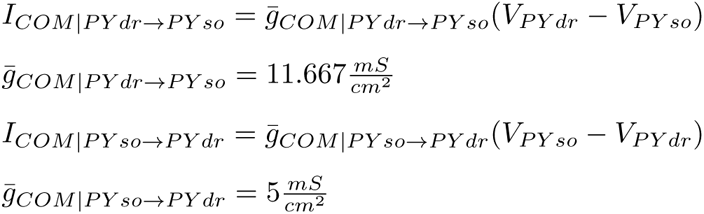

#### 7.2 (K(Na) Intercompartmental Na- activated K Channel

Additionally, there is a special and very important current, *I_K_*_(_*_Na_*_)_, which requires information on the state of the Na present in directly-connected PYso and PYdr compartments. This changes depending on ACh dose. Practically speaking, this can be modeled somewhat easily by programming it as if it is a synaptic current. The equations follow:

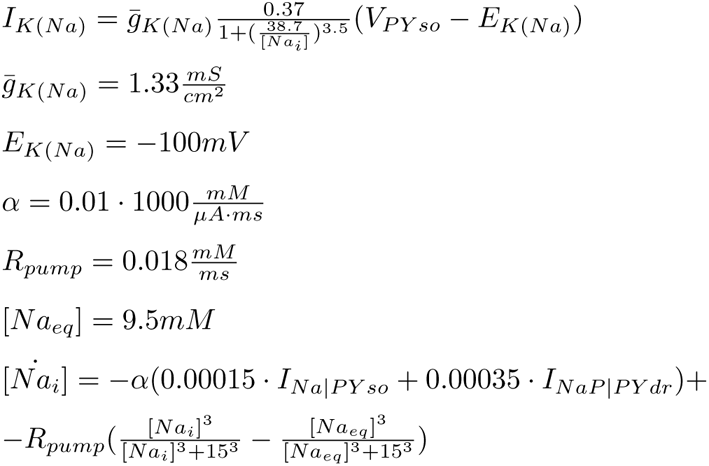

### **8** Synaptic Equations

#### 8.1 Channel Equations

The electrical current equations for each class of synapse (*AMPA*, *AMPA_D_*, *NMDA*, *GABA_A_*, and *GABA_B_*) were the same within their class, with the sole exception of different connections having different maximal conductances. All conductances of individual synapses were divided by a normalizing factor of how many connections each target cell received. Note that *AMPA_D_*is referred to as *AMPA* in the main manuscript; its different name is only relevant here for explaining that it contains depression mechanisms in the implementation.

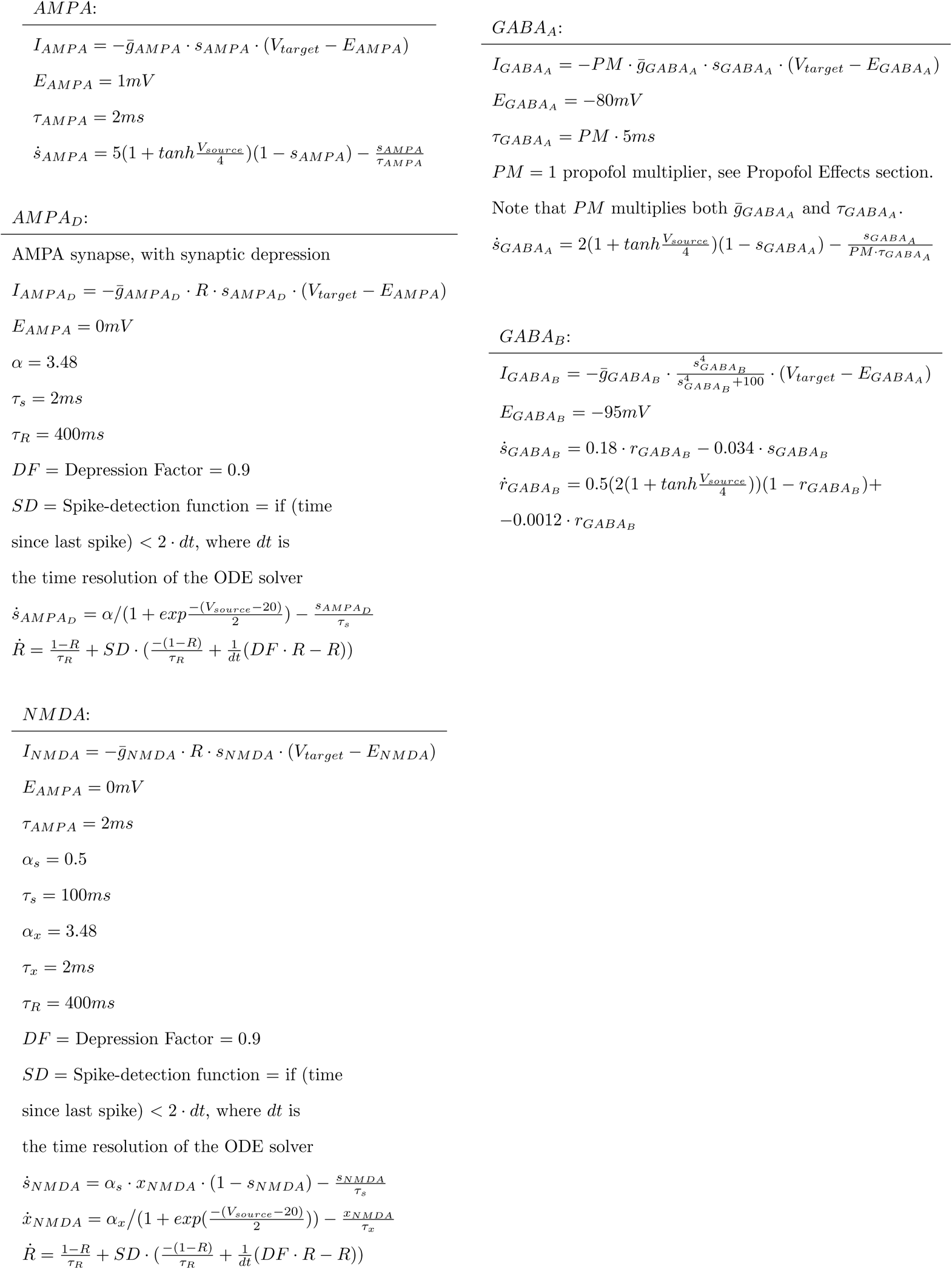

#### 8.2 Synaptic connectivity and conductances

See the Methods section for general description of connectivity. For the “nearest neighbors” connection algorithm, see the code file “models/netconNearestNeighbors.m”. Additionally, incoming synapse maximal conductances are divided by a Normalization Factor (NF). This NF ensures that the total maximal conductance for a particular synaptic type into a target cell is balanced across the number of incoming cell synapses. NF is defined as:

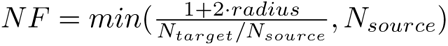

Where *N_source_*and *N_target_*are the number of cells in the source and target populations, respectively.

The following table lists all synaptic connection radii and synaptic maximal conductances (before NF has been applied). *PY so → PY dr* synaptic connections only connect to 2*·radius* target cells so as not to synapse onto their corresponding compartment. All maximal conductances are giving in units of 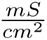. For all synapses modulated by propofol, see the following section Propofol Effects.

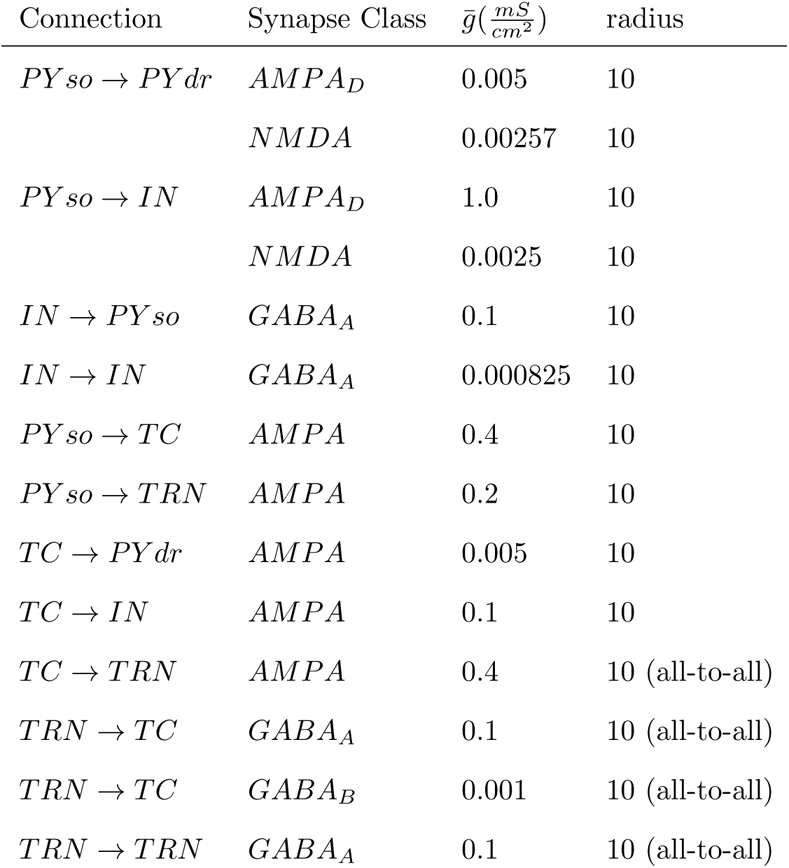

### 9 Propofol Effects

For the “wakelike” non-propofol state, see the Methods. For the anesthetic simulations, the following parameters were set. Note that PM or “Propofol Multiplier” multiplies both 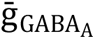 and *τGABAA*.

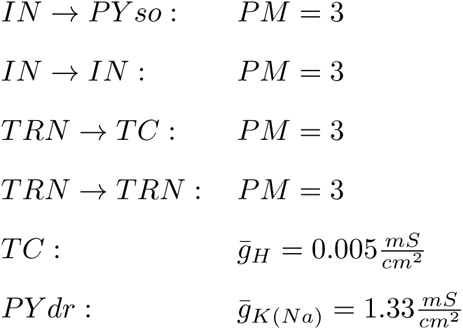

### 10 Reproducibility and Code

All simulations were run using the MATLAB-based DynaSim software package [Sherfey et al., 2018] on the “dev” branch, using MATLAB 2021a. The individual mechanism files for use with DynaSim are available online [Soplata, 2023a], and so are all the runscripts needed to reproduce simulations of the paper [Soplata, 2023b].

